# Novel native serum peptidomics workflow enables the discovery of circulating subtype-specific peptide biomarkers in acute ischemic and haemorrhagic stroke

**DOI:** 10.64898/2026.07.26.740776

**Authors:** Sachin Kote, Jakub Faktor, Marc Müller, Artur Pirog, Paulina Czaplewska, Bartosz Karaszewski, Ted Hupp, Natalia-Marek Trzonkowska

## Abstract

Novel serum peptidomics offers a direct insight into proteolytic activity, tissue injury, and systemic signaling. Nevertheless, existing workflows suffer from low peptide yields, low throughput, and limited recovery of low-abundance species. Here we present a native serum peptidomics protocol that integrates mild acid treatment, solid-phase extraction with molecular weight cutoff filtration and data-independent acquisition mass spectrometry (DIA-MS). The protocol requires less than 100 µl of serum or plasma, is completed within hours, time-cost-effective and compatible with 96-well formats without specialized equipment. Applied to a proof-of-concept cohort of patients with acute ischemic stroke (AIS), intracranial haemorrhage (ICH), and healthy controls, the workflow identified over 12,000 peptides, exceeding the three-fold threshold of existing peptidomics approaches. DIA-MS analysis across independent batches demonstrated 78–83% peptide overlap and consistent fold-change directionality. We further introduce peptide locus analysis, which aggregates overlapping peptides within defined protein regions. This approach revealed bidirectional regulation within individual precursor proteins such as the fibrinogen alpha chain (FIBA), resolving intraprotein proteolytic dynamics. Three candidate peptides from TYB4, CO4B, and ITIH4 proteins accurately distinguished stroke subtypes and controls, while characteristic shifts in peptide physicochemical properties were observed across strokes. This workflow substantially advances the sensitivity, throughput, and biological resolution of serum peptidomics for quantitative multi-biomarker discovery, validation and its output promises effective implementation of AI/ML models aiming for new dimensions in diagnostics, prognostics, prediction and monitoring.

## Introduction

Stroke is a leading cause of death and long-term disability worldwide, with acute ischemic stroke (AIS) and intracranial haemorrhagic stroke (ICH) representing mechanistically distinct subtypes that require fundamentally different clinical management [1]. Rapid and accurate subtype differentiation is essential, as thrombolytic therapy with recombinant tissue plasminogen activator (rtPA) is contraindicated in ICH and must be administered within a narrow therapeutic window in AIS [2]. Neuroimaging remains the gold standard for diagnosis; however, access to Computed Tomography (CT) or Magnetic Resonance Imaging (MRI) is not universally available, and imaging alone does not capture the molecular cascade underlying injury [3]. Blood-based biomarkers that reflect stroke-specific pathophysiology could therefore complement imaging, support treatment decisions, and enable point-of-care diagnostics.

The serum peptidome, the ensemble of endogenous peptides naturally present in circulation, is shaped by protease activity, cellular secretion, and the shedding of protein fragments during disease events, tissue injury and remodeling. Unlike tryptic peptides generated by enzymatic digestion in proteomics workflows, native serum peptides carry biological information about active proteolytic events, receptor signaling, and compartmental leakage in real time [4]. Peptidomics, the systematic identification and quantification of these endogenous peptides, therefore, offers a complementary and potentially superior layer of molecular information for biomarker discovery. Despite this promise, serum peptidomics has been technically constrained. Published workflows typically involve protein precipitation with acetonitrile or methanol, depletion of high-abundance proteins, desalting, and in some cases immunoprecipitation or enzymatic digestion. These multi-step protocols are time-consuming (3 to 5 days), costly (€60 to 80 per sample), and prone to analyte loss, particularly for low-abundance peptides. As a result, most published studies report fewer than 1 to 2,000 serum peptide identifications, a depth insufficient for robust biomarker discovery across heterogeneous patient populations [5], [6], [7], [8].

Here we address these limitations by introducing a novel, simplified, high-sensitivity native serum peptidomics workflow. Our protocol omits precipitation, depletion, desalting, and digestion, instead relying on mild acid treatment to dissociate peptides from carrier proteins, followed by HLB solid-phase extraction and molecular-weight cutoff filtration shown in Figure 1. We demonstrate that this approach identifies over 12,000 serum peptides per experiment, surpassing existing methods by at least three-fold [5], [6], [7], [8]. We apply this workflow to a stroke cohort (Healthy controls, AIS patients without ICH, and AIS patients with ICH following rtPA treatment) to characterize subtype-specific serum peptide signatures. We introduced the concept of peptide “locus” analysis to resolve bidirectional regulation within individual proteins and to present an expression-guided decision-tree biomarker classifier that stratifies AIS, ICH, and healthy controls based on discriminative peptide abundances with translational potential. In addition, observed characteristic shifts in multidimensional definitions such as native peptide physicochemical properties promise the effective implementation of Artificial Intelligence/Machine Learning (AI/ML) models aiming for new dimensions in diagnostic prediction, pattern recognition and monitoring of stroke pathology.

**Figure 1.**
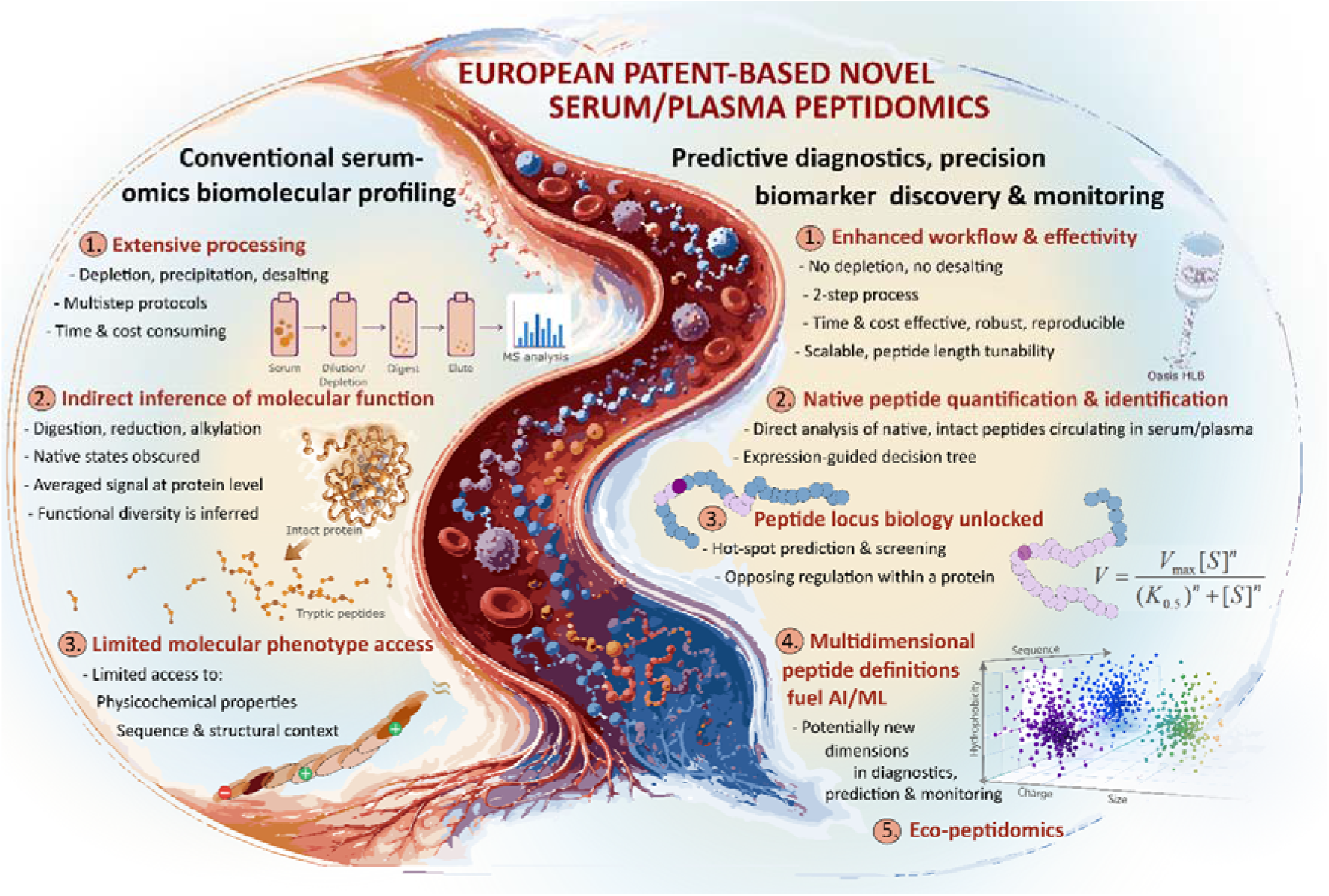
Schematic comparison of conventional serum-based omics workflows and a novel native serum/plasma peptidomics strategy. The left panel depicts standard workflows that typically involve protein precipitation, depletion, and enzymatic digestion prior to mass spectrometry, each increasing handling complexity and reducing recovery of low-abundance peptides while obscuring the native peptide context. The right panel summarises a simplified novel serum/plasma peptidomics workflow based on mild acid treatment, solid-phase extraction, and molecular weight cutoff filtration, designed to enrich endogenous circulating peptides from low serum input within hours without specialized instrumentation, followed by data-independent acquisition mass spectrometry (DIA-MS). This approach enables improved recovery, peptide coverage and quantitation of native circulating peptides. Native peptide quantitation may reveal novel biomarkers, dysregulated loci represented by overlapping peptides mapping to a defined protein region and shifts in peptide physicochemical properties for downstream biomarker modeling. It facilitates the development of global applications for human and non-human (Eco-peptidomics) sample-based diagnostics.

## Results

### Mass spectrometry data pre-processing and quality control

A project-specific spectral library was generated from DDA (data-dependent acquisition) files, including one run per serum from cohort of N = 15 subjects (5 AIS, 5 ICH and 5 healthy controls) as well as pooled fractions obtained using high-pH reversed-phase and strong cation-exchange (SCX) fractionation. The resulting spectral library comprised 12,255 peptides and was used to extract quantitative peptidomics data from serum cohort of N = 15 subjects (5 AIS, 5 ICH and 5 healthy controls). A bar plot of per sample missing values (Figure 2A) identified two outlier replicates (AIS_3 and ICH_1) with > 20% missing values and peptide intensity distributions inconsistent with other replicates (Supplementary Figure 1B). These outlier replicates were excluded from downstream quantitative analyses to minimize potential bias.

**Figure 2.**
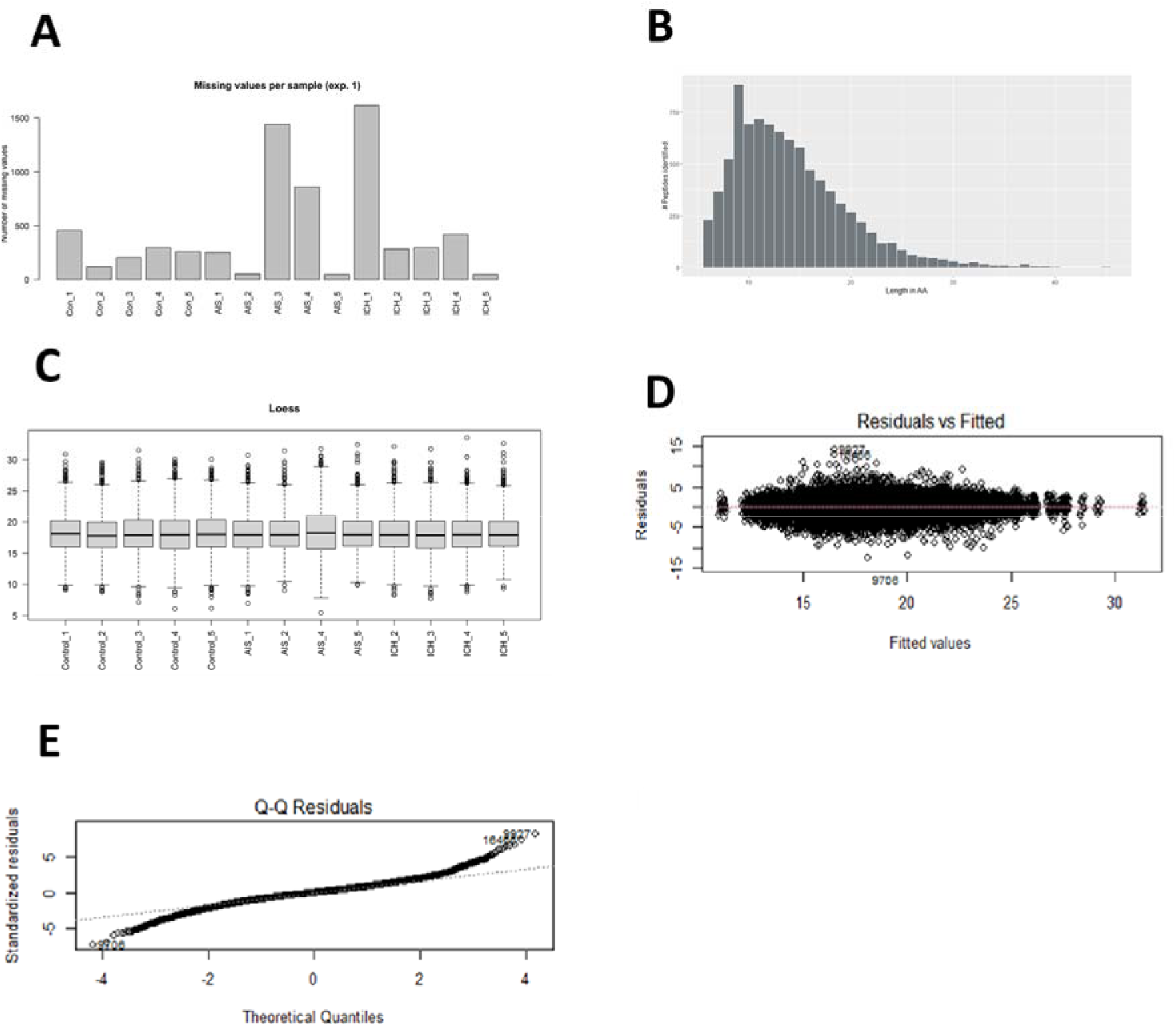
Data quality assessment and normalization performance. A) distribution of missing values across the sample set. B) peptide length distribution in the spectral library. C) Peptide intensity distributions after LOESS normalization. D) Evaluation of LOESS normalization using residuals versus fitted values. E) Quantile-quantile plot of standardized residuals showing close adherence to the reference line with mild deviation at the extreme tails, confirming effective normalization.

Following filtering, data quality was reassessed across the remaining samples (N=13, Figure 2B-E). The peptide length distribution (Figure 2B) was predominantly within the 9-40 amino acid range, consistent with typical serum peptidomics profile.

### Normalization and validation

LOESS normalization of extracted log2-transformed peptide intensities mitigated inter-sample variability. Post-normalization sample-wise, summed peptide intensity bar plots showed aligned medians and comparable interquartile ranges (Figure 2C). Normalization quality was confirmed by a symmetric residual-vs-fitted intensity plot centered at zero (Figure 2D) and a quantile-quantile plot of standardized residuals that closely tracks the 45-degree reference line, with a mild tail flaring (Figure 2E), consistent with effective normalization.

### Global peptidotype correlations and clustering

Pearson correlation analysis of quantitative peptidomics sample fingerprints (peptidotypes) revealed positive inter-sample correlation across all comparisons (Supplementary Figure 1A). Correlation coefficients (r=0.44-0.89) and regression fits indicated positive correlations with significant p-values (***: p ≤ 0.01) in all comparisons (Supplementary Figure 1A). The peptide intensity heatmap (Supplementary Figure 1B) revealed distinct clustering of control subjects from stroke patients, reflecting global differences in serum peptidomics. Control samples 1, 3, and 5 showed the highest intra-group corelations (r=0.86-0.89). In contrast, haemorrhagic and ischemic stroke peptidotypes (Supplementary Figure 1B) clustered together, but exhibited subtle profile differences.

A filtered heatmap retaining peptides with the highest fold-change standard deviation (STDEV cutoff > 3.4) (Supplementary Figure 1C) enhanced stroke subtype separation while maintaining the distinct control cluster, identifying a high-variance candidate biomarker pool with subtype-specific expression patterns. To further investigate potential biomarkers, serum peptide fold-changes between AIS, ICH and controls were calculated from quantitative DIA data.

### Benchmarking against existing serum peptidomics and proteomic workflows

The workflow was benchmarked against three closely related reported serum peptidomics methods using the same stroke sample set in two independent experiments (Exp1 and Exp2), and the results are summarized in Table 1. Table 1 shows that none of the compared methods achieved an efficiency sufficient to identify and quantify the number of peptides comparable to that of the novel method. The novel workflow identified at least ten-fold more peptides than any of the benchmarked methods, demonstrating substantially superior sensitivity for peptide detection (Table 1).

**Table 1.** Wet-lab benchmark of the novel serum peptidomics method against three concurrent methods. The combination of an HLB column and a 3 kDa filter enabled more effective and more pure isolation of the serum peptidome, yielding at least 10-fold more identified peptides than the compared methods.

| Method description | Exp1 | Exp2 | Method 1 | Method 2 | Method 3 |
| --- | --- | --- | --- | --- | --- |
| Sample input | 100 μl | 100 μl | 500 μl | 50 μl | 50 μl |
| Number of samples | 15 | 15 | 2 | 1 | 2 |
| Time required | 240min + 240min | 240min + 240min | 24 h | 24 h | 24 h |
| Filtration | HLB | HLB | No | ultrafiltration | ultrafiltration |
| Precipitation | No | No | ACN | No | No |
| Depletion | No | No | No | No | No |
| LC/MS method | 120 min/DDA | 120 min/DDA | 120 min/DDA | 120 min/DDA | 120 min/DDA |
| Average Peptides identified per sample | 7197 | 6051 | 437 | 120 | 558 |
| Qualitative analysis possible | Yes | Yes | Marginally | No | Marginally |
| QC passed using TIC | Yes | Yes | No | No | No |

Extended benchmarking through a systematic literature review (Table 2) confirmed that existing serum peptidomics methods identify up to approximately 2,000 native serum peptides, while serum proteomics workflows relying on tryptic digestion identify up to approximately 1,700 proteins. The novel workflow achieved at least three-fold more native serum peptide identifications in both independent experiments (Exp1 and Exp2; Table 2), with superior quantitative accuracy (Table 3). Crucially, the protocol achieves this without protein precipitation, desalting, depletion, or immunoprecipitation, relying solely on acid retrieval of peptides from carrier protein and HLB column solid-phase extraction followed by centrifugal 3/10 kDa cutoff filtration.

**Table 2.**
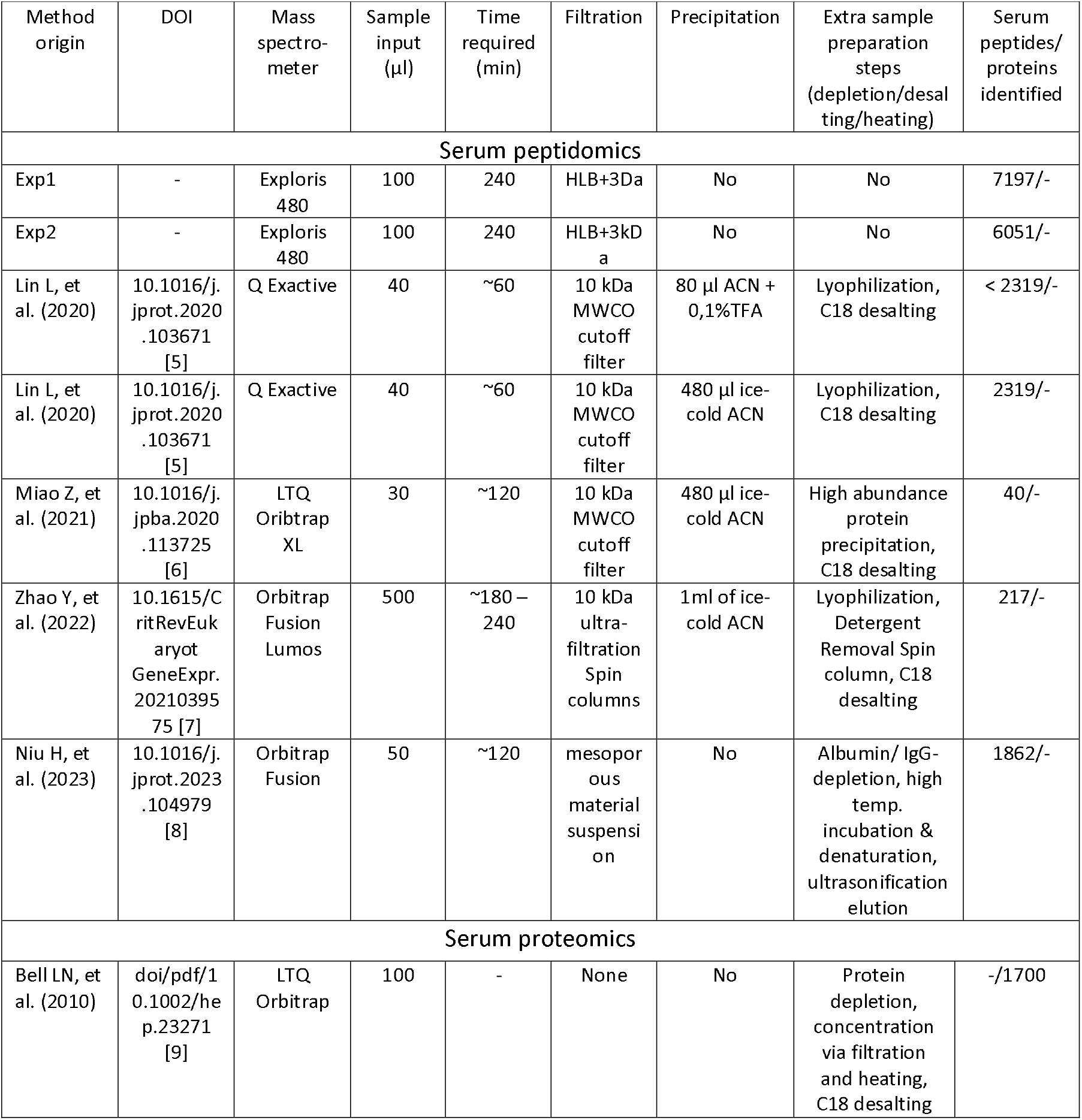
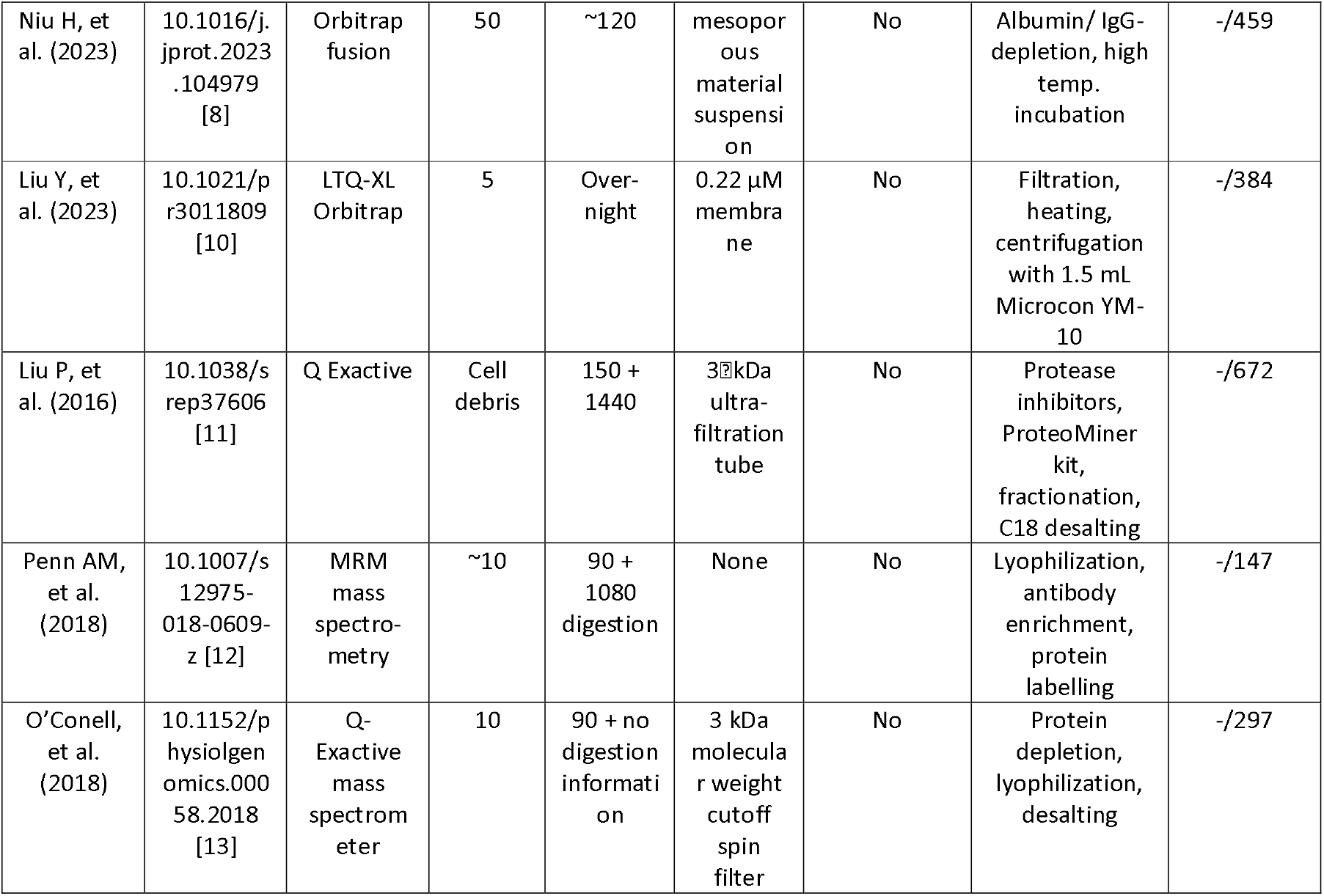
Benchmark of the new serum protocol against current serum peptidomics and proteomics workflows identified through a literature review. The table is divided into two sections. The first section includes methods focused on the analysis of native serum peptides, which are most closely aligned with our novel protocol. The second section summarizes proteomics approaches that target protein identification via enzymatic digestion, thereby obscuring information on native peptide forms. Across all evaluated methods, our protocol demonstrates at least a three-fold increase in the detection of native serum peptides and proteins.

**Table 3.**
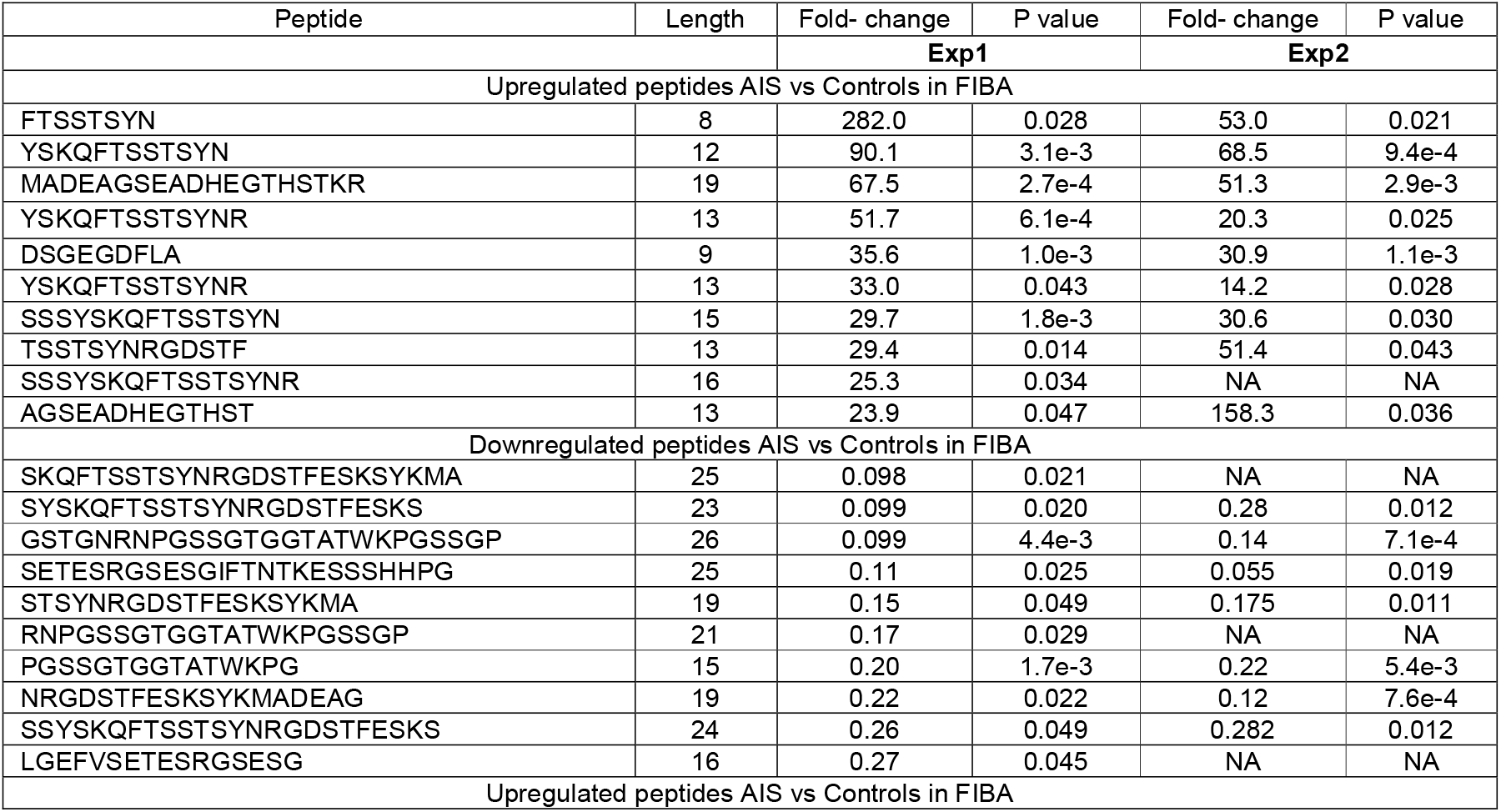
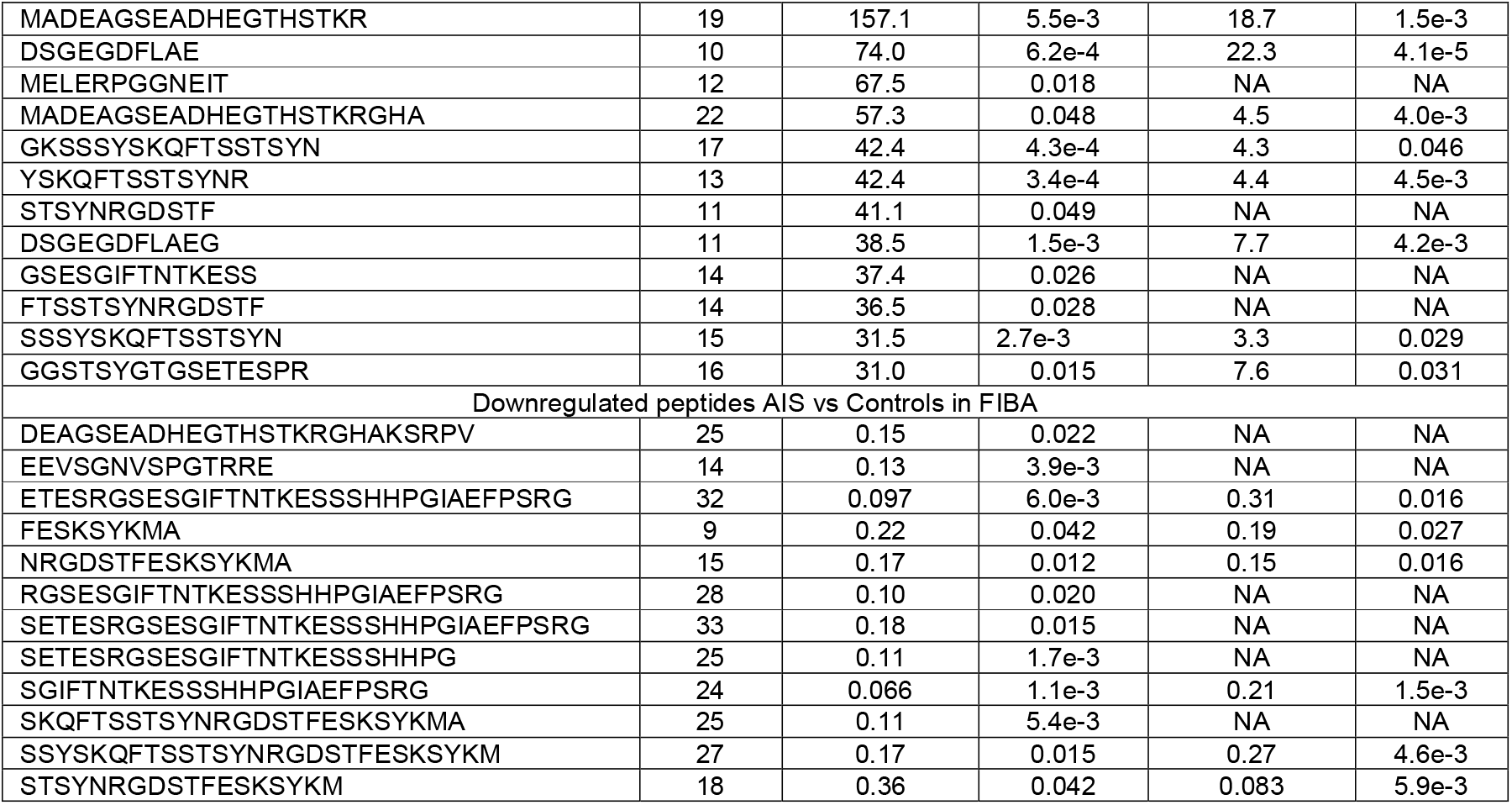
Selected peptides to evaluate fold-change reproducibility between two independent datasets (Exp1(n=13) and Exp2 (n=15)) derived from a single serum sample set processed in separate batches. Consistent fold-changes and P-values across datasets demonstrate the quantitative reproducibility of the novel serum peptidomic workflow for both down- and up-regulated peptides.

### Quantitative reproducibility across independent experimental batches

To evaluate workflow robustness, the same serum cohort (N=15) was processed in two independent experimental batches approximately one year apart (Exp1 and Exp2), incorporating minor protocol refinements while preserving the core workflow. Distinguishing serum peptidotypes between AIS and ICH requires high analytical precision and reproducibility. Spectral libraries generated in PEAKS Studio identified 12,255 peptides (FDR 1%) in Exp1 and 10,769 peptides (FDR 1%) in Exp2 (Figure 3). Across conditions, the lowest peptide count was observed in the Exp2 ICH group (5,910 peptides) and the highest in the Exp1 AIS group (6,197 peptides). Group-wise comparisons of identified peptide sequences revealed high reproducibility, with peptide sequence overlap between corresponding groups ranging from 78% to 83% (Figure 3; Supplementary Figure 2A-B). Venn diagram analyses within and across both experiments identified 2,755 peptides shared across AIS, ICH, and control groups in both Exp1 and Exp2 (Supplementary Figure 2C). Each group contained relatively few experiment-specific peptides (approximately 200-800 per condition), further supporting high inter-experiment concordance.

**Figure 3.**
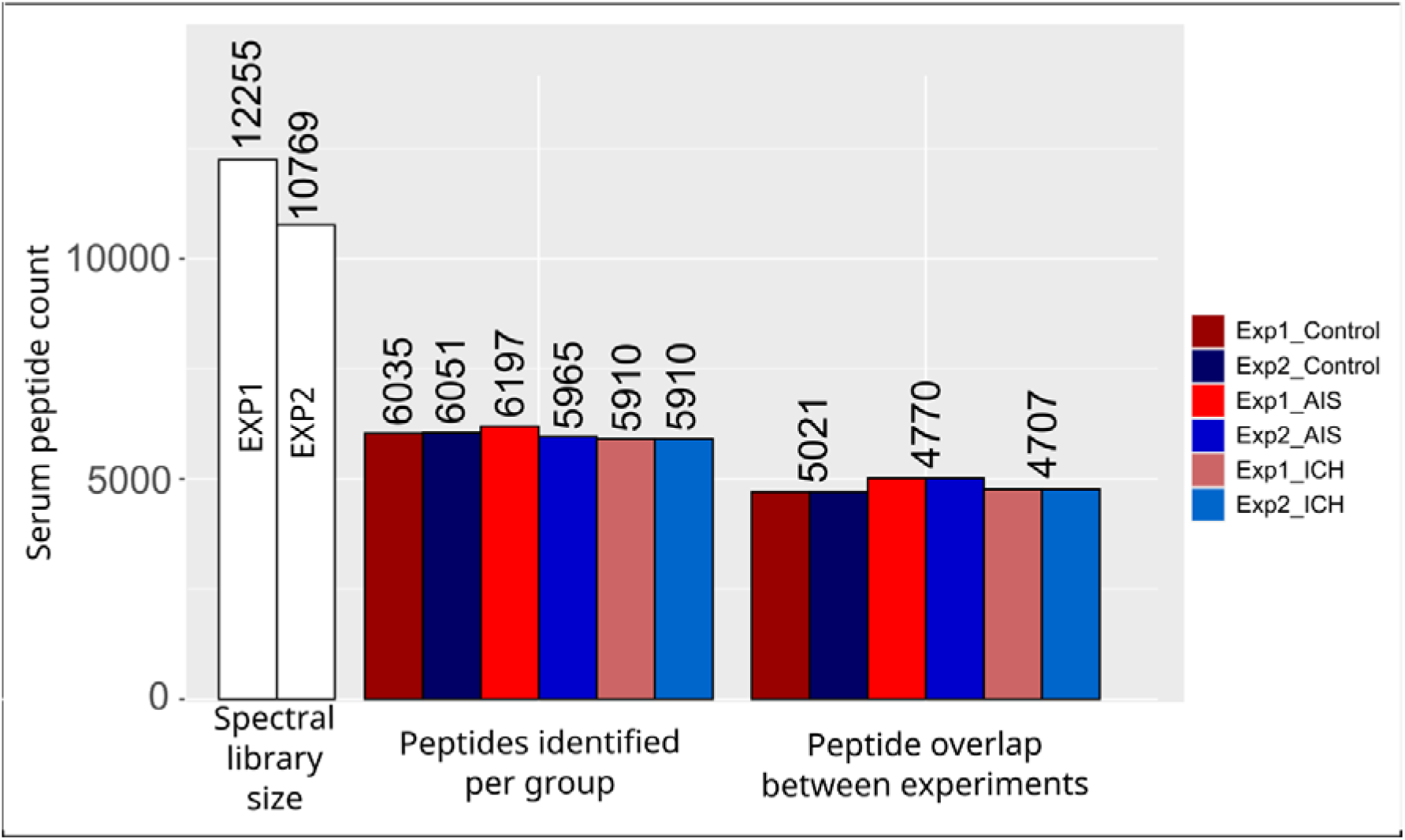
Comparison of spectral libraries and groups across experimental batches. Overlap of identified serum peptide sequences between Exp1 and Exp2 for AIS (acute ischemic stroke), ICH (intracranial haemorrhagic stroke) and control groups. The high proportion of shared peptides across corresponding groups demonstrates the reproducibility of the peptidomics workflow.

Quantitative reproducibility was assessed using the most up- and down-regulated peptides derived from fibrinogen alpha chain (P02671, FIBA_Human) and additional top-dysregulated peptides filtered from full quantitative analysis shown in Supplementary Table 1 (Table 3). Across all comparisons, peptide fold-changes in Exp1 and Exp2 followed consistent directional trends when detected in both datasets (Table 3). Together, Figure 3, Table 3 and Supplementary Figure 2 A-C demonstrate accurate peptide sequence identification and consistent quantitative performance, supporting the suitability of the protocol for biomarker discovery in large serum cohorts.

### Revealing biologically significant differential peptide regulation in acute ischemic and haemorrhagic stroke

Follow-up quantitative peptidomic analysis was restricted to 2488 serum peptides quantified in at least 3 samples per condition, and with the quantification FDR was less than 1%. The analysis revealed significant peptide dysregulation across all pairwise comparisons of strokes and controls. The number of significantly up- and down-regulated peptides are summarised in Table 4, with corresponding sequences and fold-changes traceable in full quantitative results in Supplementary Table 1. Differential peptide regulation is visualized by volcano plots in Supplementary Figures 3-5, which visualize the -log_10_ adj. p-value (cutoff = 0.05) against log_2_ of its fold-change (cutoff > 0.58 and <-0.58). Supplementary Figure 3 visualizes peptide dysregulations in acute ischemic stroke (AIS) compared to control, Supplementary Figure 4 visualizes peptide dysregulations in intracranial haemorrhagic stroke (ICH) compared to control, and Supplementary Figure 5 directly compares both stroke subtypes.

**Table 4.** Numbers of up- and down-regulated serum peptides identified in quantitative comparisons among control, intercranial haemorrhagic stroke (ICH) and acute ischemic stroke (AIS). Peptides were considered significant at an adjusted P-value ≤ 0.05 and a log_2_ fold-change ≥ 0.58 or ≤ −0.58.

| Comparison | N up-regulated peptides (adj.pval sig.) | N down-regulated peptides (adj.pval sig.) | No significant regulation |
| --- | --- | --- | --- |
| Both strokes to control | 348 | 171 | 1969 |
| AIS to control | 211 | 123 | 2154 |
| ICH to control | 337 | 164 | 1987 |
| ICH to AIS | 88 | 59 | 2341 |

Follow-up quantitative peptidomic analysis was restricted to 2488 serum peptides quantified in at least 3 samples per condition, and the quantification FDR was less than 1%. Volcano plot analysis of serum peptidome identified a subset of peptides with significant abundance changes between stroke and control samples, indicating peptide-level alterations related to stroke biology. Due to the limited availability of stroke peptidomics mass spectrometry datasets for direct benchmarking, we mapped these dysregulated peptides to their protein precursors, revealing signatures associated with known stroke-related biological processes. Including inflammation and complement activation, endothelial injury, extracellular matrix (ECM) degradation, lipid metabolism alterations, oxidative stress, neurorepair, cytosolic leakage from injured cells, damage-associated molecular pattern (DAMP) release, platelet activation, and microthrombosis (Supplementary Table 2) with further details discussed later.

### Expression-guided decision-tree classification of stroke subtypes

The novel serum peptidomic data (Supplementary Table 1, 2) together with existing literature, suggest that both ischemic and haemorrhagic stroke affect sets of native peptides differing primarily in the magnitude and temporal dynamics of these changes with biological relevance. To identify the most discriminative peptides for stroke subtype classification, an expression guided decision-tree analysis was applied to the quantified peptide pool (Figure 4A). This decision tree showcases high-value biomarker candidates selected from the broader quantified peptide pool. The first node of the decision tree (Figure 4A) distinguishes healthy controls from stroke patients based on the NPLPSKETIEQEKQAGES (Thymosin beta-4) peptide intensity (Kruskal-Wallis, p-value = 1.69E-06). The second node uses intensity cutoffs of the (CO4B) DAPLQPV peptide to discriminate intracranial haemorrhage stroke (ICH) patients from acute ischemic stroke (AIS) patients (Kruskal-Wallis, p-value = 0.0003). Finally, intensity cutoffs on SSRQLGLPGPPDVPDHAAYHPF (Inter alpha-trypsin inhibitor, heavy chain 4/ITIH4) peptide were used to further identify and confirm acute ischemic stroke (Kruskal-Wallis, p value = 3.9E-05).

**Figure 4.**
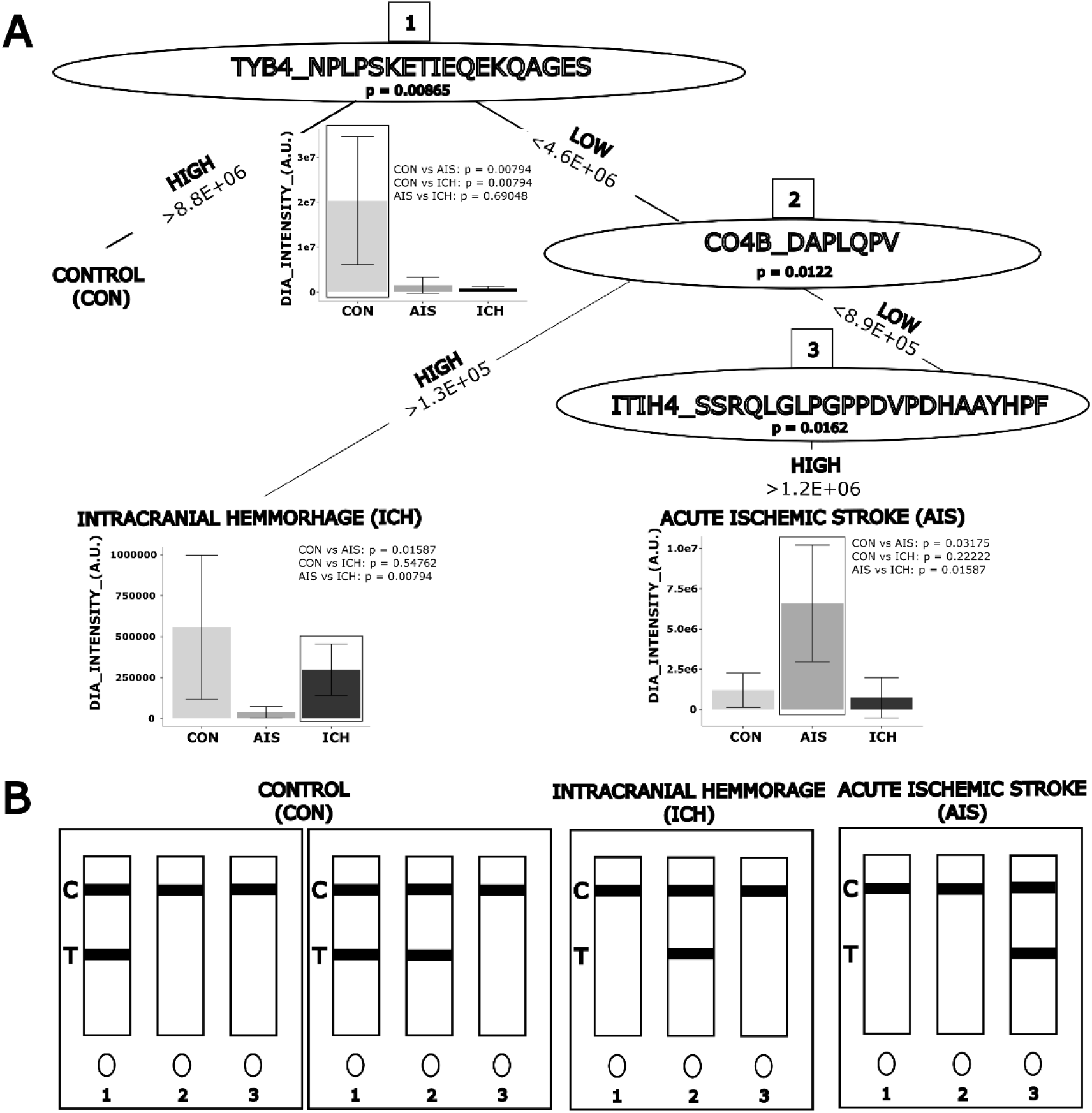
Translating DIA-based serum peptidomics into an expression-guided decision tree and theoretical lateral flow assay for stroke classification. A) Expression-guided decision tree stratifying controls (CON, N=5), acute ischemic stroke (AIS, N=4) and intracranial haemorrhagic stroke (ICH, N=4) based on DIA-MS intensities of selected serum peptide biomarkers. Kruskal-Wallis P-values are indicated within decision nodes, and Mann-Whitney P-values for pairwise comparisons are shown alongside the corresponding bar plots. Bar plots represent mean peptide intensities with error bars indicating standard deviation across subjects. B) Conceptual three-way peptide-based lateral flow assay illustrating the potential clinical translation of the identified biomarkers for stroke subtype classification.

To illustrate the potential clinical translation of these biomarkers, a conceptual three-way lateral flow assay prototype incorporating these peptides is presented in Figure 4B, demonstrating how DIA-based peptide intensities could be implemented in a simple, rapid, and cost-effective point-of-care diagnostic format upon clinical validation.

### Physicochemical feature comparison between AIS and ICH dysregulated peptide sets

A preselection step was applied to identify peptides up- and down-regulated in intracerebral haemorrhage (ICH) relative to acute ischemic stroke (AIS), as well as in AIS- and ICH-associated peptides relative to controls. These subsets were subjected to *in-silico* physicochemical characterization, including molecular weight, peptide length, isoelectric point, net charge, charge density, and hydrophobicity-related metrics. Features showing the strongest differences between ICH and AIS populations were then identified.

As expected from the preselection strategy, fold-change displayed complete separation between the groups (KS = 1.00, Cohen’s d = 1.60). Among the physicochemical properties, the largest differences were observed for molecular weight (KS = 0.61, d = 1.37), charge density (KS = 0.56, Cohen’s d = 1.27), isoelectric point (KS = 0.55, Cohen’s d = 1.12), net charge (KS = 0.53, Cohen’s = 1.21), and peptide length (KS = 0.48, Cohen’s d = 1.09). Moderate separation was also observed for aliphatic index and instability-related descriptors (Figure 5). Distinct physicochemical profiles between the two stroke subtypes (Figure 5) indicate that peptide populations associated with AIS and ICH differ not only in abundance but also in their physicochemical composition.

**Figure 5.**
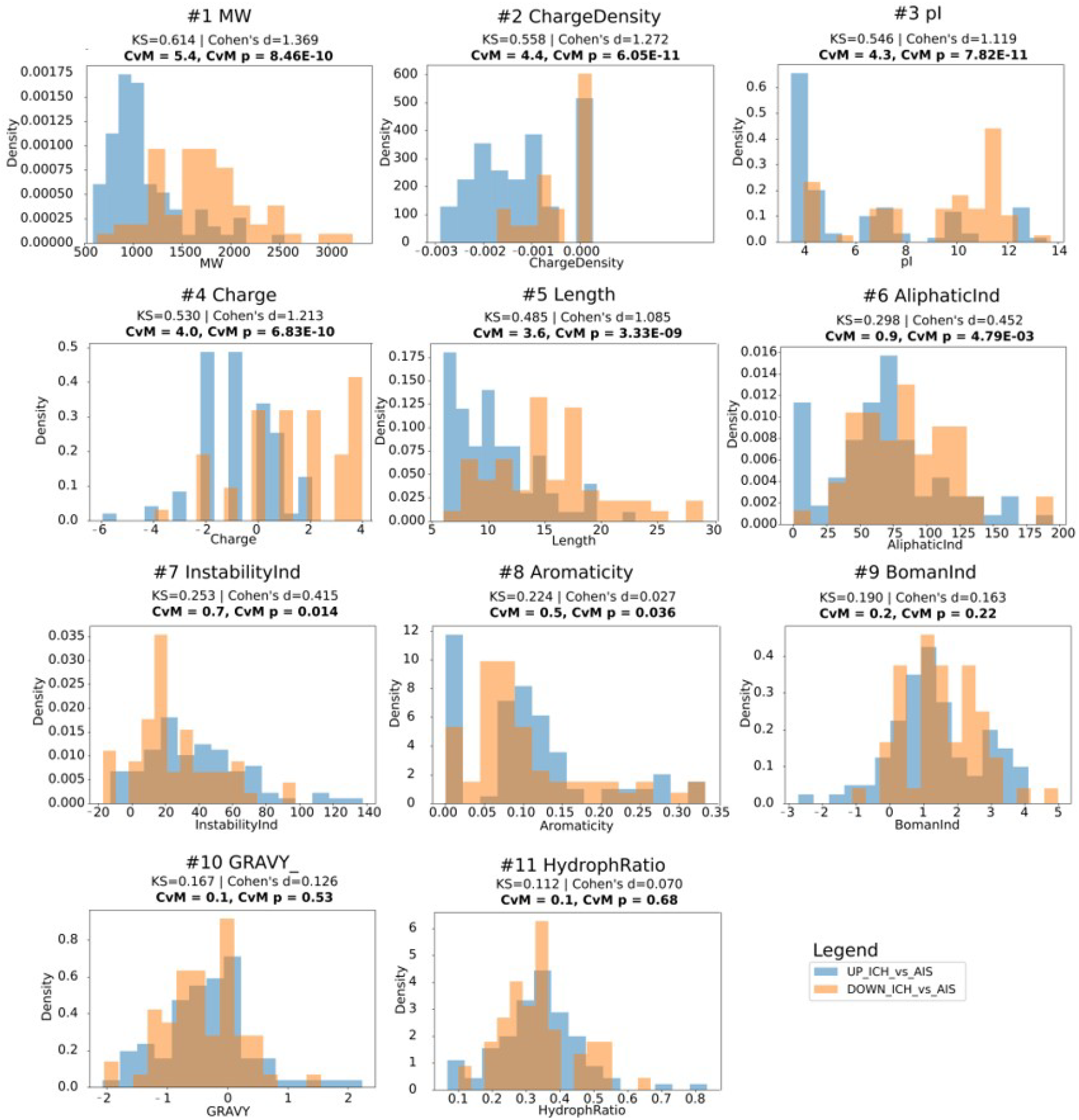
Distributional comparison of physicochemical properties of peptides up-regulated and down-regulated in the acute ischemic stroke (AIS) and intracerebral hemorrhage (ICH) comparison. Histograms show the distributions of physicochemical properties calculated for peptide subsets identified as up- and down-regulated following differential abundance analysis between AIS and ICH. Differences between peptide populations were quantified using mean shifts, Cohen’s *d* effect sizes, two-sample Kolmogorov–Smirnov (KS) statistics, and pairwise two-sample Cramér–von Mises (CvM) statistics, which were subsequently used to rank physicochemical properties according to the degree of distributional separation. Histograms are ordered in descending order of KS and CvM statistics, with molecular weight exhibiting the strongest separation between peptide subsets, followed by charge density, isoelectric point, net charge, and peptide length. Physicochemical properties displaying lower KS values showed progressively weaker discrimination, with hydrophobicity-related features exhibiting the smallest distributional differences. The observed shifts indicate that peptide populations associated with AIS and ICH differ not only in abundance but also in their underlying physicochemical properties.

Further comparisons were made to assess whether these physicochemical alterations will also be evident in control subjects (CON) compared with AIS- and ICH-associated peptide populations. In both comparisons, molecular weight showed the strongest shift, accompanied by consistent differences in peptide length. Relative to controls, stroke-associated peptides exhibited lower molecular weights and shorter lengths, with more pronounced effects in ICH (MW: Cohen’s d=0.90; length: Cohen’s d=0.80) than in AIS (MW: Cohen’s d=0.65; length: Cohen’s d=0.60). A shift toward shorter, lower-molecular-weight peptides was observed in both conditions (Supplementary Figure 6), consistent with enhanced proteolytic fragmentation initiated in stroke. Beyond size-related features, differences were also detected in aromaticity, instability-related descriptors, and charge-associated properties. Notably, AIS showed stronger alterations in charge-related features, including net charge, charge density, and isoelectric point, suggesting enrichment of peptides with distinct electrostatic characteristics (Supplementary Figure 6). Overall, stroke-associated peptide populations differed substantially from controls, with the most prominent changes involving peptide size and charge-related properties.

### Native peptide locus (loci) analysis reveals stroke-specific regulation within a single protein sequence

To exploit peptidomics analysis beyond protein-level measurements, we introduced a peptide locus (loci) as a cluster of overlapping endogenous peptides mapping to the same region of a precursor protein sequence (including isoforms) and evaluated for coordinated regulation. This approach was applied to fibrinogen alpha chain (FIBA; P02671) which overall showed highest overall sequence coverage by quantified native peptides (Tables 5 and 6; Figure 6).

**Figure 6.**
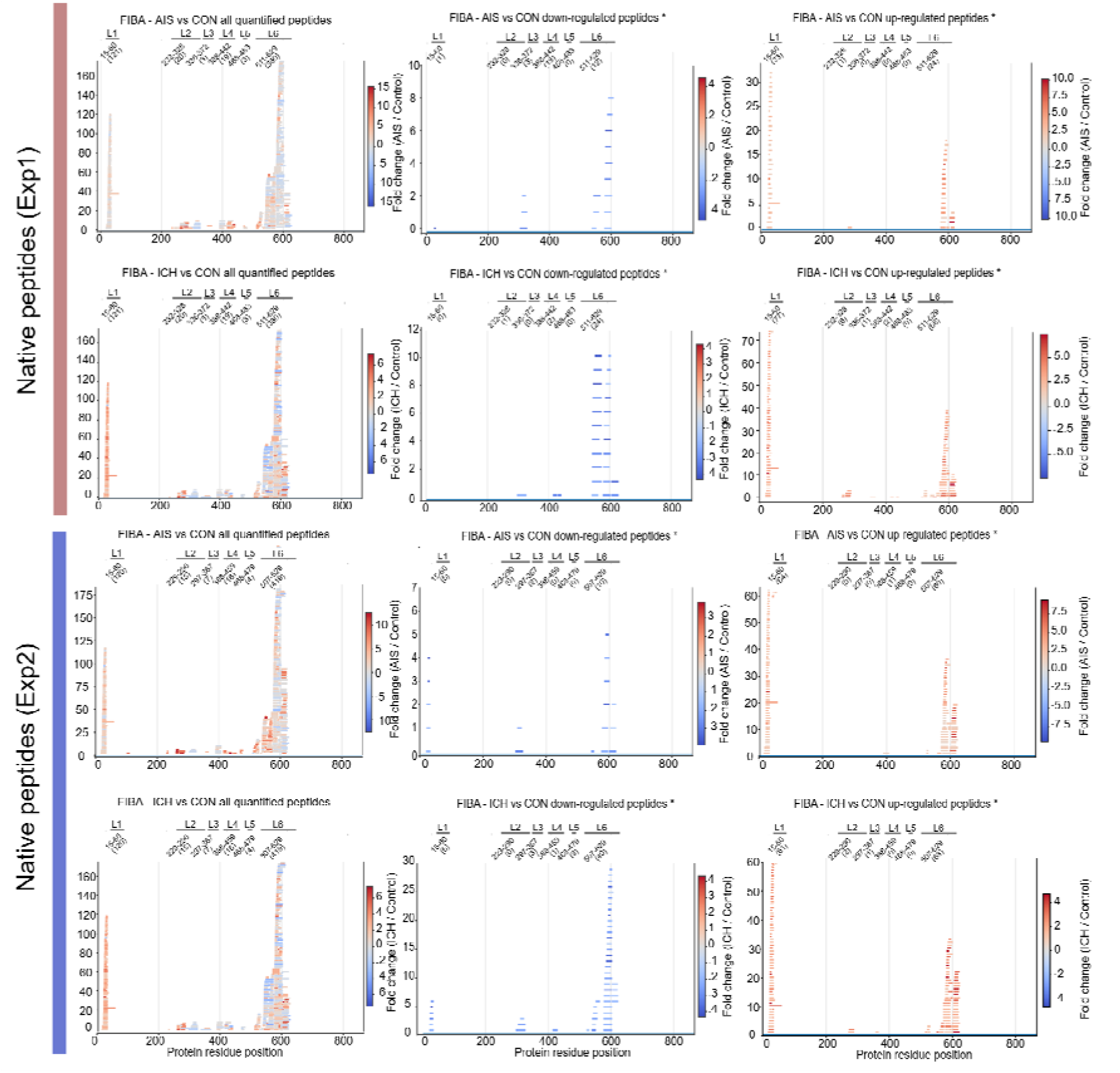
FIBA peptide loci were evaluated in acute ischemic stroke (AIS) vs controls (CON) and intracerebral haemorrhage (ICH) vs CON. Locus 1 was uniquely upregulated in strokes vs CON, supported by consistent up-regulation of multiple peptides, In contrast, loci 6 is exhibited mixed regulation, limiting their discriminatory value at the locus position level, however, it is slightly apparent that there is different density and length of up- and down-regulated native peptides. Moreover, the differences in density and length of up- and down-regulated peptides within locus 6 discern the ICH and AIS. These results highlight the importance of peptide-level analysis beyond protein-level measurements for biomarker discovery. Moreover, native peptide loci occupation lays foundation for AI/ML pattern recognition to distinguish ICH and AIS, emphasizing subtype-specific biomarker potential. *Significantly dysregulated peptides (adjusted P-value ≤ 0.05 and a log_2_ fold-change ≥ 0.58 or ≤ −0.58).

**Table 5.** Top dysregulated native serum peptide counts per protein and their regulation across ICH vs control, AIS vs control and ICH vs AIS comparisons. *We are not referring to unique peptides that would resolve/distinguish between the isoforms. **canonical isoform refers to the canonical Uniprot entry. ^#^adjusted P-value ≤ 0.05 and a log_2_ fold-change ≥ 0.58 or ≤ −0.58.

| Comparison | Protein/Isoform | N up-regulated peptides<br>(adj.pval sig.)<br>(Exp1)<sup>#</sup> | N down-regulated peptides<br>(adj.pval sig.)<br>(Exp1)<sup>#</sup> |
| --- | --- | --- | --- |
| AIS vs control | P02671/FIBA canonical isoform | 58 | 16 |
|  | P02671/FIBA isoform 1 | 34 | 0 |
|  | P02671/FIBA isoform 3 | 33 | 0 |
|  | P02671/FIBA isoform 5 | 33 | 0 |
|  | Q14624/ITIH4 canonical isoform | 27 | 0 |
|  | Q14624/ITIH4 heavy chain 4 isoform | 24 | 0 |
|  | Q14624/ITIH4 unannotated isoform | 20 | 0 |
|  | P60709/ACTB canonical isoform | 0 | 15 |
|  | P63261/ACTG canonical isoform | 0 | 14 |
| ICH vs control | P02671/FIBA canonical isoform | 154 | 0 |
|  | P02671/FIBA isoform 1 | 77 | 0 |
|  | P02671/FIBA isoform 3 | 84 | 0 |
|  | P02671/FIBA isoform 5 | 77 | 0 |
|  | P50221/MEOX2 canonical isoform | 76 | 0 |
|  | P02671/ITIH4 canonical isoform | 21 | 0 |
|  | SRGN canonical isoform | 10 | 0 |
| ICH vs AIS | P02671/FIBA canonical isoform | 36 | 16 |
|  | P02671/FIBA isoform 1 | 31 | 0 |
|  | P02671/FIBA isoform 5 | 30 | 0 |
|  | alpha fold protein of fibrinogen alpha | 30 | 0 |
|  | P02675/FIBB canonical isoform | 9 | 0 |
|  | P50222/MEOX2 canonical isoform | 30 | 0 |

**Table 6.** Identified peptide loci within FIBA sequence harbouring overlapping native serum peptides.

| Experiment |  | Locus | StartAA - lastAA | N quantified native peptides | N significantly down-regulated native peptides | N significantly up-regulated native peptides | Comment |
| --- | --- | --- | --- | --- | --- | --- | --- |
| Experiment 1 (Exp1) | AIS vs CON | 1 | G15-W60 | 121 | 1 | 33 | Stroke-specific locus |
|  |  | 2 | L232-P328 | 20 | 0 | 1 | Non-specific locus |
|  |  | 3 | S335-H371 | 3 | 3 | 0 | Non-specific locus |
|  |  | 4 | S388-L442 | 19 | 0 | 0 | Non-specific locus |
|  |  | 5 | T468-V483 | 3 | 0 | 0 | Non-specific locus |
|  |  | 6 | H511-V629 | 380 | 12 | 24 | Different density of native peptide occupancy among strokes, different length of up- and down-regulated peptides |
|  | ICH vs CON | 1 | G15-W60 | 121 | 0 | 77 | Stroke-specific locus |
|  |  | 2 | L232-P328 | 20 | 1 | 6 | Non-specific locus |
|  |  | 3 | S335-H371 | 3 | 0 | 1 | Non-specific locus |
|  |  | 4 | S388-L442 | 19 | 2 | 2 | Non-specific locus |
|  |  | 5 | T468-V483 | 3 | 0 | 0 | Non-specific locus |
|  |  | 6 | H511-V629 | 380 | 24 | 68 | Different density of native peptide occupancy among strokes, different length of up- and down-regulated peptides |
| Experiment 2 (Exp2) | AIS vs CON | 1 | G15-W60 | 120 | 5 | 64 | Stroke-specific locus |
|  |  | 2 | V229-S290 | 15 | 0 | 0 | Non-specific locus |
|  |  | 3 | S297-R367 | 7 | 2 | 0 | Non-specific locus |
|  |  | 4 | S388-R459 | 16 | 0 | 1 | Non-specific locus |
|  |  | 5 | T468-T479 | 4 | 0 | 0 | Non-specific locus |
|  |  | 6 | D507-V629 | 419 | 10 | 60 | Different density of native peptide occupancy among strokes, different length of up- and down-regulated peptides |
|  | ICH vs CON | 1 | G15-W60 | 120 | 6 | 61 | Stroke-specific locus |
|  |  | 2 | V229-S290 | 15 | 0 | 2 | Non-specific locus |
|  |  | 3 | S297-R367 | 7 | 3 | 1 | Non-specific locus |
|  |  | 4 | S388-R459 | 16 | 1 | 0 | Non-specific locus |
|  |  | 5 | T468-T479 | 4 | 3 | 0 | Non-specific locus |
|  |  | 6 | D507-V629 | 419 | 40 | 68 | Different density of native peptide occupancy among strokes, different length of up- and down-regulated peptides |

Across the dataset, protein-level comparisons masked peptide-level regulation, with simultaneous peptide up- and down-regulation observed within single protein sequences, most prominently in FIBA. Within FIBA, multiple loci composed of overlapping peptides showed significant but bidirectional regulation across comparisons (Table 5, 6). Supporting this, the analysis was performed across two independent experiments (Exp1 and Exp2) to assess the reproducibility of locus generation, with each experiment quantifying 546 and 581 native FIBA peptides mapping into six major peptide loci (Figure 6).

Loci 1 and 6 showed the most pronounced differences between control and stroke samples. Locus 1 (G15–W60) was predominantly occupied by peptides exhibiting increased abundance in stroke samples, with significantly up-regulated peptides comprising 27–64% of all quantified peptides within this locus depending on stroke type and experiment, while significantly down-regulated peptides remained minimal (0–5%). This highly asymmetric regulation pattern, was consistent across experiments, supporting locus 1 as a robust stroke-associated region. Locus 6 (H511-V629/D507-V629) displayed a more complex distribution, containing both up-regulated and down-regulated peptides, however, the overall density of peptide occupancy increased markedly in stroke samples, with the most pronounced enrichment observed in ICH, where significantly up-regulated peptides reached 16.8% of the locus total peptides in Exp1 and 16.2% in Exp2 which is up to two-fold higher than in AIS at the same locus (Exp1 6.3% and Exp2 14.3%) (Figure 5). Further separation of peptides according to their regulation direction within locus 6 revealed differences in peptide characteristics between stroke subtypes. Up-regulated peptides within this locus were represented by a higher number of shorter peptides, whereas down-regulated peptides were less abundant but displayed a tendency toward longer peptide sequences which is partly in line with observations from physicochemical properties analyses in Figure 5. The combination of high peptide density, heterogeneous regulation patterns, and reproducible locus organization highlights locus 6 as a complex source of native peptide information that may contain individual biomarker candidates associated with stroke pathology. This pattern might suggest differential intensity of proteolytic processing of the FIBA between stroke subtypes.

Loci 2-4 showed substantially lower peptide occupancy and regulation rates; however, their distribution suggested a preferential representation in ICH compared with AIS. However, with current dataset no consistent directional signal was observed across experiments and therefore these loci were classified as non-specific. The observed differences in locus composition, peptide density, regulation direction, peptide length, and cross-experiment reproducibility demonstrate that native peptide loci capture multiple layers of molecular information associated with stroke status and subtype. Locus 1 emerges as the strongest candidate for a general stroke indicator. Loci are functionally important sites of peptide release in stroke pathophysiology.

### Biological relevance investigation through native peptide inference to corresponding proteins

Both ischemic and haemorrhagic strokes trigger early biochemical events which include endothelial stress, ECM breakdown and collapse of cytoskeleton induced by either hypoxia-mediated ischemia or mechanical vessel rupture [14]. Our peptidomics data reveals dysregulation of cytoskeletal and cytosolic proteins including actins [15], PDLIM1/PDLIM5 [16], [17], vimentin [18], [19], [20] and septin2 [21], consistent with endothelial disruption followed by vascular leakage. Metabolic stress is evident from dysregulated ATP-citrate synthase [22], [23], PYGB [24], [25], adenylyl cyclase associated protein [26], [27] and flavin reductase (NADPH) fragments [28], [29], reflecting damage to neuronal, glial, endothelial and recruited immune cells, potentially leading to vascular leakage. Dysregulation of peptides from endothelial ECM proteins such as fibrillin-1 [30], [31], LTBP2 [32], MGP [33], ITIH2/ITIH4 [34], [35], [36], [37] and MMRN1 [38] in both ischemic and haemorrhagic stroke, as reflected by altered levels of peptide precursors in our assay (Supplementary Table 2), indicates proteolytic degradation of ECM, expectedly more prominent in ICH due to physical vascular injury but also triggered in AIS likely via reperfusion-activated proteases.

Detected dysregulation of haptoglobin [39], [40], hemopexin [41], [42], [43], and ceruloplasmin [44] indicates perturbed hemoprotein processing, and oxidative iron homeostasis in stroke pathology. This dysregulation evident in both ICH and AIS, likely reflects blood spill out from vessel rupture or oxidative injury during reperfusion and suggests a shared iron-induced oxidative stress and hemoprotein scavenging across stroke subtypes [45]. Nuclear proteins such as Histone H1.2 [46], dysregulated in stroke patient serum, represent damage-associated molecular patterns (DAMPs) characteristic for the transition from the tissue injury to inflammatory activation. Markers of significant complement activation and inflammation were also detected [47] including dysregulation of complement components such as complement factor B (CFAB) [46], complement C3 [48], C1S [49], C1QC [50], [51], C4B [52], CFAD [53], [54] which contribute to neutrophil recruitment [55], tissue damage, vascular occlusion and blood-brain barrier disruption. Our serum peptidomics assay revealed related dysregulation of peptides from inflammation-associated mediators: such as SAA1 [56], Seglycin [57], CCL/CXCL7 [58], [59], highlighting their role in post-stroke inflammation and tissue remodeling. Elevated protease inhibitors including serpins (SERPINA1, SERPINA3, kalistatin (SERPINA4)) [60], Galectin-1 [61], [62], [63] and alpha-2-macroglobulin (A2MG) [64], [65], suggest an onset of counter-regulatory response limiting tissue destruction. Notably, complement driven blood-brain disruption enabled detection of central nervous system derived-proteins such as IGF-axis components, vimentin [18], [66], actins [15], [67] and DAMPs in serum. Dysregulation of multiple apolipoproteins such as APOA [68], APOB [69], [70], [71], APOC [72], APOF [73], [74], APOA4 [75], APOA2 [76], [77], APOE [77] further indicates modulation of innate immunity, lipoprotein remodeling and repair pathways post stroke. Proteins associated with neuronal repair and angiogenesis such as, such as QSOX1 [78] and IGF2 [79], [80] were also altered, suggesting an early tissue repair. Moreover peptides linked to coagulation and platelet activation pathways CXCL7 [58], [59], FYB1 [81], [82], MMRN1 [83], Septin2 [84], BIN2 [85] and factors involved in clot formation, fibrinolysis and vascular permeability such as fibrinogen [86], prothrombin [87], coagulation factors [88], KNG1 [89] and KLKB1 [90], [91] were detected, reflecting the distinctly triggered but overlapping mechanisms of AIS and ICH (Supplementary Table 2).

### Broad applicability across diseases and biofluid types

The novel serum peptidomics workflow was applied to additional patient cohorts beyond stroke to demonstrate its generalisability (Table 7). Experiments on cohorts with cardiovascular disease, autoimmune conditions (including type 1 diabetes), non-small cell lung cancer (NSCLC), and oesophageal adenocarcinoma (OAC) all yielded high-quality MS-compatible samples with robust peptide identifications (Table 7). Independent technical replicates performed on the same serum sets confirmed protocol reproducibility. These results indicate that the workflow is broadly applicable across human (serum/plasma) disease contexts and is potentially extendable to non-human biological samples. Moreover, it yields robust, enhanced peptide identification relative to established peptidomics workflows (Table 1, 2, 7) and integrates diverse bioinformatic pipelines. Currently, the workflow is successfully extended to other biofluids, such as urine and synovial fluid.

**Table 7.** Overview of serum peptidomics experiments performed using novel method on different diseases and patient cohorts through different data analysis pipelines. *Technical replicates, **Two independent experiments on same sera set.

| Disease | Sub-condition | Cohort (N) | MS runs* (N) | Serum peptides quantified | Pipeline used |
| --- | --- | --- | --- | --- | --- |
| Cardiovascular/<br>stroke | Acute ischemic stroke | 5 | 25 | 4329/3384** | PEAKS studio |
|  | Haemorrhagic stroke | 5 | 25 |  |  |
|  | Healthy control | 5 | 25 |  |  |
| Autoimmune<br>/Diabetes<br>mellitus type 1 | Standard treatment | 7/7 | 21/21 | 8439 | MS-Fragger,<br>Skyline,<br>MSstats |
|  | Treg treatment | 8/8 | 24/24 |  |  |
|  | Treg + mab treatment | 6/6 | 18/18 |  |  |
|  | Patients at admission | 7 | 21/21 |  |  |
|  | Healthy control | 9 | 27 |  |  |
| Non-small cell<br>lung cancer | Cancer patients | 9 | 27 | 8600 | DIA-NN,<br>Perseus |
|  | Healthy controls | 9 | 27 |  |  |
| Oesophageal<br>carcinoma | Cancer patients | 4 | 12 | 2122 | MS-Fragger,<br>Skyline,<br>MSstats |
|  | Healthy controls | 4 | 12 |  |  |

## Discussion

We present a novel serum peptidomics method that enhances the sensitivity, simplicity, and biological resolution of native peptide mass spectrometry in complex biofluids such as blood. By combining mild acid extraction at pH 3.1-3.6 with HLB solid-phase enrichment and molecular-weight cutoff filtration, the method eliminates the precipitation, depletion, digestion and desalting steps that have traditionally limited throughput and peptide recovery in serum/plasma peptidomics/proteomics. The workflow is scalable, 96-well compatible, and processes <100 µl of serum/plasma per sample within hours at a reagent cost of approximately €5-7, compared with 3-5 days and €60-80 for conventional approaches. The molecular-weight cutoff step is tuneable, with the 10 kDa configuration used here representing one implementation within a broader 3-30 kDa range, and the protocol also supports optional on-filter retentate enzymatic digestion. Together, these features preserve a wide native peptide range while enabling practical high-throughput implementation for serum/plasma and other biofluids.

The impact of our purification strategy is reflected in the improved native peptide recovery. Conventional acetonitrile precipitation denatures carrier proteins but can co-precipitate peptide complexes, trapping low-abundance analytes in insoluble aggregates and reducing native peptide signal. In contrast, mild acid incubation dissociates native peptides from carrier proteins while preserving their solubility. The subsequent Oasis HLB extraction step recovers native peptides while retaining larger proteins and matrix components, with partial desalting. Final molecular-weight cutoff filtration further polishes the peptidome by removing residual high-molecular-weight contaminants, enriching native peptides in the flow-through. The combined effect of reduced handling and milder extraction yields >12,000 native serum/plasma peptides, representing a several-fold increase over current state-of-the-art and our initial trial protocols.

Beyond analytical depth, the method showed robust biological reproducibility. Quantitative analysis yielded sample-specific peptide fingerprints and introduced the peptidotype, analogous to the established proteotype. Peptidotype correlations across technical replicates were strong. In biological replicates, correlations improved after standard missing-value filtering, and stroke samples separated cleanly from controls in unsupervised analyses. This robustness was confirmed by an independent experiment performed ∼1 year later with an unchanged core workflow. Within-group peptide sequence overlap was ≥78% and fold-change directions were consistent across experiments, indicating that these signals are reproducible features of the serum peptidome.

The reproducibility of the data suggests that serum peptidomics can capture both individual biomarker changes and the global remodeling of endogenous peptide generation driven by biochemical processes involved in stroke. The biological meaning of individual dysregulated peptides was first inferred from the corresponding proteins. Up-regulation of peptides derived from cytoskeletal and endothelial structural proteins point to vascular disruption and cellular leakage, while ECM-associated peptides indicate matrix remodeling. Coordination in processes of inflammation, oxidative stress, coagulation activation, and endogenous protective anti-inflammatory responses documented by a shift in peptide signals from haemoprotein scavenging proteins, complement components, acute-phase mediators, and their counter-regulatory inhibitors further reveals result’s biological significance.

Importantly, although many dysregulated peptides and molecular events overlap between AIS and ICH, their context differ, supporting stroke subtype-specific diagnostics rather than a single generic injury signature (Supplementary Table 2). The peptidomic biomarker signatures observed here are therefore biologically grounded and reflect core stroke injury mechanisms rather than nonspecific systemic noise [92], [93], [94]. Beyond canonical fold-change-driven peptidomic shifts, stroke pathology appears to induce coordinated shifts in molecular weight, peptide length, net charge, charge density, and isoelectric point indicating that stroke-associated peptide populations occupy distinct regions of physicochemical space. This implies that AIS and ICH are not defined solely by isolated peptide abundance changes, but by broader disease-specific peptidomic features, consistent with enhanced proteolytic fragmentation initiated in stroke. Shifts across this dense molecular feature space may support machine-learning (AI-ML)-based approaches for disease-specific natural peptide pattern recognition, improved prognostics, diagnostics, predictions, and monitoring performance.

A central analytical contribution of the study is the identification of the most discriminating candidate native peptide biomarkers for AIS and ICH, using both differential expressions ranking and a peptide locus framework. The native peptide loci provide a perspective on disease-induced proteolytic changes that is not captured by protein-centric analysis. The FIBA sequence reveals this with particular clarity given its role in haemostasis, platelet degranulation, and fibrin clot formation [86], likely reflecting specific proteolytic events induced by vascular injury and thrombosis. The six loci identified across two independent experiments map onto the known functional architecture of FIBA with remarkable precision. Locus 1 (G15–W60) within the N-terminal region represents thrombin cleaved fibrinopeptide A, released upon fibrin polymerization trigger [95]. Its dominant reproducible up-regulation in both AIS and ICH is consistent with thrombin-driven coagulation activation at the stroke site with a greater magnitude observed in ICH, potentially reflecting more diffuse coagulation engagement accompanying vascular rupture [96]. It should be noted that serum collection involves *in vitro* coagulation, which generates fibrinogen-derived peptides shared across all sample groups. Therefore, the stroke-associated fold-changes reported here represent differential signals above this uniform baseline rather than a preanalytical bias. Locus 6 (D507–V629) maps onto the αC domain, which undergoes conformational unfolding upon fibrinopeptide B cleavage to expose binding sites for plasminogen, making it the target for fibrinolytic machinery and a proteolytic hotspot [97]. The increased peptide density at locus 6, most pronounced in haemorrhagic stroke, may reflect heightened exposure of circulating fibrinogen to activated coagulation pathways and proteolytic enzymes, while at slightly lesser extent analogous processes likely occur in AIS due to vascular injury and coagulation activation. The simultaneous presence of up- and down-regulated peptides within locus 6 likely further highlights the complexity of stroke-associated proteolysis, or alternatively an artifact of sample processing and/or measurement. The pattern of shorter up-regulated peptides and fewer, longer down-regulated peptides in locus 6 may reflect altered proteolysis or clot incorporation, but the mechanism remains unclear.

Routine proteomics sample preparation includes the depletion of highly abundant proteins, and FIBA is the most common highly abundant protein depleted from samples for downstream biomarker discovery. In contrast, the novel peptidomics emphasize the biological role of natural peptides from high-abundant proteins such as FIBA. The spatial organization of peptide signals across FIBA supports a biologically coherent observation confirming that the observed serum peptidomic signatures reflect structured *in vivo* processes such as proteolysis, rather than nonspecific degradation. Together, native peptide loci represent molecular patterns integrating peptide localization, abundance, sequence length, and directionality of regulation. These findings establish a strong rationale for integrating multi-protein peptide loci into future AI/ML diagnostic frameworks, where biologically structured signals may substantially enhance classification accuracy beyond single-protein models. Such a multidimensional signature provides a natural, biochemically driven basis for ML/AI classification approaches, potentially enabling better discrimination of biological states than current state-of-the-art methods.

Translationally, the differential expression guided decision tree illustrates how DIA-derived intensities of discrete native peptides can be converted into interpretable classification rules. Using a composite ranking of effect size and statistical significance, we identified a compact panel of peptides, including sequences (NPLPSKETIEQEKQAGES) derived from Thymosin beta-4, CO4B (DAPLQPV), and ITIH4 (SSRQLGLPGPPDVPDHAAYHPF), that stratified controls, AIS, and ICH in a proof-of-concept cohort. This expression-guided decision tree must be regarded as exploratory, since feature pre-selection and model construction occurred within the same cohort and the sample size was limited. Nevertheless, the fact that the top-ranked natural peptides map to proteins involved in inflammatory activation, complement biology, and cytoskeletal remodeling supports the biological coherence of the discriminatory signal and argues that compact peptide panels may have future diagnostic utility [34], [36], [98].

Taken together, these findings position DIA-based serum/plasma peptidomics as a scalable platform for systems-level stroke biomarker discovery. The workflow combines practical advantages in speed, cost, and sample compatibility with improved analytical depth and reproducibility, enabling native peptide level resolution that supports both biomarker discovery and mechanistic interpretation. By preserving native peptides, exposing locus-specific features, and capturing stable peptidotype structure, the method creates a framework for studying disease as a coordinated peptidotype fingerprint rather than as a collection of isolated marker changes. Future studies in larger, independent cohorts will be required to validate the proposed peptide panels and decision rules. However, this work establishes a foundation for clinically interpretable, high-resolution serum peptidomics in acute cerebrovascular disease.

## Limitations

Several limitations of the present study merit consideration. 1) We emphasize that the current study is a discovery-phase analysis in a small, well-characterized cohort (N = 15). 2) The sample size of five subjects per group limits statistical power and generalisability. 3) Validation in larger, independent cohorts with diverse clinical presentations, comorbidities, and sampling time points is required before clinical implementation. 4) Longitudinal studies examining peptidome dynamics across the acute, ICH, and recovery phases of stroke will be essential to define the diagnostic window and assess prognostic utility. 5) The heterogeneity inherent to serum peptidomes influenced by age, sex, lifestyle, and comorbidities not controlled for here likely contributes to the partial co-clustering of AIS and ICH samples. 6) Additionally, the functional significance of the identified peptide loci requires experimental validation, including assessment of whether specific peptides are causally related to stroke pathophysiology or are secondary epiphenomena.

### Future directions include

i) prospective validation of candidate peptide biomarkers in large, multi-centre stroke cohorts; (ii) integration of the peptidomics workflow with orthogonal validation platforms (ELISA, targeted mass spectrometry, lateral flow assay); (iii) extension to longitudinal sampling to capture the temporal evolution of the serum peptidome post-stroke; and (iv) application of the peptide locus framework to other complex diseases where protein-level biomarker discovery has failed or haled. v) Ongoing study to show the impact of sample collection, handling, serum/plasma types, types of anticoagulants, with/without proteases, etc., on peptidomes. vi) Disease-specific peptide locus will be emerging as “hot spots”. With AI-pipelines, these peptide “hot spots” facilitate the screening and development of early diagnostics, monitoring, prediction, prognosis, and therapy development for acute as well as chronic diseases.

In summary, the novel native serum peptidomics workflow presented here represents a significant methodological advance, offering unique peptide depth, high reproducibility, and biologically coherent discovery performance. The peptide locus concept and characteristic shifts in multidimensional definitions such as native peptide physicochemical properties provides a new analytical lens for resolving the complexity of the serum peptidome, and the identified stroke biomarker candidates offer a promising foundation for the development of rapid, multiplexed diagnostic tools. Platform well-suited to multi-biomarker discovery in stroke and, potentially, in other acute conditions where rapid, blood-based diagnosis is clinically valuable.

## Materials and methods

### Study participants and ethics

Blood samples were collected from patients admitted to the University Clinic in Gdansk in collaboration with Prof. Bartosz Karaszewski, head of the Department of Neurology at the Medical University of Gdansk. This study was conducted in accordance with the principles of the Declaration of Helsinki (as revised in 2024). All participants provided written informed consent prior to inclusion in the study. The study was approved by the Independent Bioethics Commission at the Medical University of Gdansk. A total of 15 serum samples (n=15), each derived from a distinct individual, were included in this study. Samples were divided into 3 groups (n=5 per group): i) healthy controls (CON), ii) patients with acute ischemic stroke treated with recombinant tissue plasminogen activator (rtPA) without subsequent intercranial haemorrhage (AIS), and (iii) patients with AIS treated with rtPA who developed intercranial haemorrhage (ICH).

Each sample was prepared individually according to the following protocol below and afterwards stored at -80°C for possible further usage. Peripheral venous blood was drawn to Becton Dickinson Vacutainer tubes containing clot activator. Samples were collected on the day of inclusion into the study. For the rtPA treated patients, blood collection was performed after the treatment administration. Following collection, blood samples were allowed to clot at room temperature for up to 40 min. Serum was then separated by centrifugation at 1000-2000 x g for 10 min at room temperature. The resulting supernatant was transferred to low-binding microfuge tubes and initially stored at -20°C for several hours prior to long-term storage at -80°C.

All samples were processed according to standard clinical protocols implemented at the Neurological Department of the Medical University of Gdansk. Subsequently, samples were transported under controlled cold-chain conditions to the Intercollegiate Faculty of Biotechnology of the University of Gdansk and then stored at -80 °C. To preserve sample integrity, all specimens were processed promptly after collection, subjected to timely centrifugation, handled in low-binding microfuge tubes, and maintained at controlled temperatures with minimal exposure to room temperature throughout handling and storage.

### Isolation of native serum peptides from patient sera

All buffers were prepared on the same day as the peptidomics sample preparation. A citrate phosphate buffer at pH 3.1-3.6 was prepared and adjusted to pH 3.1–3.6 using 1 M NaOH prepared in LC-MS grade water. For serum peptide extraction, 100 μL of serum from each sample was incubated with 30× citrate phosphate buffer for 5 min on ice. Samples were gently vortexed to dissociate protein-peptide interactions, then centrifuged at 5,000×g for 5 min at 4°C. The supernatant was collected and diluted 1:1 with 0.2% FA in LC-MS grade water (*v/v*), yielding a final volume of approximately 5-7 ml per sample.

For peptide purification, all diluted samples were subjected to Oasis hydrophilic-lipophilic balanced (HLB) cartridges (30 mg; Waters). Cartridges were first conditioned with 0.2% FA/methanol (*v/v*) per cartridge, then equilibrated with 0.2% FA/water (*v/v*) per cartridge. A total of 5-7 ml of diluted sample was loaded onto each cartridge. Cartridges were washed three times with water/5% methanol/0.2% FA (*v/v*) per cartridge per wash. Bound peptides were eluted with water/80% methanol/0.2% FA (*v/v*) per cartridge, and the eluate was subsequently diluted to water/40% methanol/0.2% FA (*v/v*).

Purified serum peptide samples were then processed through 3 kiloDalton (3 kDa) molecular weight cutoff (MWCO) ultrafiltration devices (Amicon Ultra-2 Centrifugal Filters, Ultracel-3K, 2 ml; Millipore) to eliminate abundant proteins. Purification steps are not restricted to one size of molecular weight cutoff (MWCO) ultrafiltration; 3/10/30 kDa devices can be used. Each 3 kDa ultrafiltration device was rinsed with 40% methanol in LC-MS-grade water (*v/v*) per column, then centrifuged at 4,000×g for 30 min at 4°C. The rinsed device was inserted into a new collecting tube, and approximately 2 ml of eluted and diluted peptide solution was pipetted into the device, ensuring the pipette tip did not touch the membrane. The device was centrifuged at 4,000×g for 135 min at 4°C. The filtrate containing purified peptides was collected and lyophilized to dryness. All dried peptide samples were stored at −80°C until mass spectrometry analysis. Serum peptides from each patient were measured three times: one data-dependent acquisition (DDA) run, and two data-independent acquisition (DIA-MS) runs.

### Native serum peptide pre-fractionation

A pooled sample was created by mixing 2 μl from each serum/plasma peptidomics sample from the proof-of-concept patient cohort. Two orthogonal fractionation methods were employed: high-pH fractionation and strong cation exchange (SCX) fractionation. For high-pH fractionation, we used the Pierce High pH Reversed-Phase Peptide Fractionation Kit (P/N 87777, Thermo Scientific). Native peptide prefractionation was performed according to the manufacturer’s instructions. Pooled serum samples were additionally fractionated by SCX. In-house C18-SCX StageTips were prepared as described in Rappsilber et al [99]. A C_18_ disk was stacked on top of a SCX disk inside 200 μl pipette tip. TFA-based SCX buffers followed protocol described in Adachi et al [100]. Fractions were collected in separate low-binding microfuge tubes and evaporated, using a SpeedVac vacuum centrifuge. The samples were then evaporated and stored at -80°C until analysis on the Exploris480 MS. Prior LC-MS/MS analysis, the samples were dissolved in 30 μl of loading buffer and the concentrations were checked at 220 and 280 nm using a Nanoready F-1100 spectrophotometer (LifeReal). Each fraction was 2x injected and analysed in data-dependent mode (DDA) on Exploris 480 (Thermo Scientific).

### Liquid Chromatography separation (LC) prior tandem mass spectrometry (MS/MS)

The eluted and dried serum peptide samples were resuspended in 30 μl of loading buffer composed of 0.08% trifluoroacetic acid (TFA) in water with 2.5% acetonitrile (ACN). Indexed retention time (iRT) peptides (P/N Ki-3002-2, Biognosys) were added according to the manufacturer’s guidelines. Then 6 μl of the dissolved sample were injected into UltiMateTM 3000 RSLCnano liquid chromatograph (Thermo Scientific) online coupled with Orbitrap Exploris 480 mass spectrometer (MS) (Thermo Scientific). μ-precolumn C18 trap cartridges (300 μm i.d.) and 5 mm length packed with C18 PepMap100 sorbent with PepMap 5 μm sorbent (P/N: 160454, Thermo Fisher Scientific) were used to concentrate and desalt serum peptides using a 5 μl/min flow of loading buffer. Peptides were then eluted on 75 μm ID and 150 mm length fused-silica analytical column packed with PepMap 2 μm sorbent (P/N: 164534, Thermo Scientific). Analytical peptide separation was performed by a non-linear increase of a mobile phase B (0.1% FA in ACN) in a mobile phase A (0.1% FA in water). A non-linear gradient started at 2.5% B linearly increasing up to 35% B in 80 min, followed by a linear increase up to 60% B in the next 15 min with a flow rate of 300 nl/min. Native serum peptides eluting from the column were ionized in a nano-electrospray ion source and were introduced to Exploris 480. LC separation parameters were kept identical across MS acquisition methods.

### Parameters of data-dependent and data-independent acquisition

Exploris 480 acquired the data in data-dependent peptide mode. The full scan was operated in profile mode with 120000 resolution, scanning the precursor range from m/z 350 Th to m/z 1650 Th. Normalized AGC (Automatic Gain Control) target was set to 300% with 100 msec maximum injection time, and each full scan was followed by fragmentation and acquisition of the top 20 the most intense precursor ions if their MS/MS spectra were not listed in dynamic exclusion list. Dynamic exclusion list contained precursors for 20 sec after their first fragmentation. Precursor isotopologues were excluded from experiment, and mass tolerance was set to 10 ppm. Minimum precursor ion intensity was set to 3.0e3, and only precursor charge states of +1 to +6 were included in the experiment. The precursor isolation window was set to 1.6 Th. Normalized collision energy type with fixed collision energy mode was selected. The collision energy was set to 30%. Orbitrap resolution was set to 60000. Normalized AGC target was set to 100% with an automatic setting of maximum injection time, and data type was centroid.

During DIA analysis Orbitrap Exploris 480 mass spectrometer operated in positive polarity and each DIA cycle was accompanied by one full-scan in profile mode with 60000 resolution. The full-scan range was set from m/z 350 Th up to m/z 1450 Th, and the normalized AGC target was set to 300% with 100 msec maximum injection time. Each DIA cycle was accompanied by the acquisition of 62 precursor windows/scan events. DIA precursor range was set from m/z 350 Th up to m/z 1100 Th with 12 Th window width and 1 Th window overlap. Normalized collision energy type with fixed collision energy mode was selected to fragment precursors included within each isolation window. The collision energy was set to 30% and Orbitrap resolution in DIA mode was set to 30000. Normalized AGC target was set to 1000% with automatic setting of maximum injection time while data type was profile.

## MS Data analysis

### Spectral library creation and quantitative data extraction

A project-specific spectral library was generated from data-dependent acquisition (DDA) measurements obtained from individual serum/plasma samples included in the proof-of-concept cohort. To maximize peptide representation, DDA datasets generated from both strong cation exchange (SCX) and reversed-phase fractionation strategies were incorporated into library construction. Raw spectral quality was initially assessed to ensure data integrity prior to database searching.

Spectral identification was performed using PEAKS Studio™ 11 against the Homo sapiens Swiss-Prot/TrEMBL database (2024) supplemented with indexed retention time (iRT) peptide sequences (Biognosys), reversed decoy sequences, and common contaminant proteins. Search parameters included a non-specific enzyme search with precursor mass tolerance of 12 ppm and fragment ion tolerance of 0.02 Da. Variable modifications included N-terminal acetylation, methylation, and methionine oxidation.

The resulting spectral library was subsequently applied for targeted extraction and quantification of endogenous serum/plasma peptides from data-independent acquisition (DIA) datasets acquired in technical duplicates for each participant. Peptide intensities were extracted using the quantitative analysis PEAKS Studio™ 11 module and normalized based on internal indexed retention time (iRT) peptide standards to correct for potential analytical variation between measurements. Peptide quantification was performed using both spectral library-assisted extraction and database search approaches using the parameters described above with the following filtering criteria: fold-change range ≥1 and ≤64, average peptide intensity ≥200, and peptide quality score ≥20. Statistical significance was assessed using analysis of variance (ANOVA), and signal normalization was performed using total ion current (TIC) normalization.

### Quantitative data analysis

Processed quantitative data were exported for downstream analysis in R Studio 4.4.2. Data quality assessment, normalization, data handling, statistical analysis, and visualization were performed using the R packages limma 3.60.4 [101], tidyverse 2.0.0 [102], ggplot2 3.5.2, and Plotly 4.11.0.

For peptide annotation and locus-based analysis, identified peptide sequences were mapped to their corresponding precursor proteins based on PEAKS Studio™ protein assignments. Protein accession identifiers were reviewed to remove redundant entries and annotated using UniProt database information.

Differential abundance analysis was performed to identify peptides and proteins exhibiting significant changes between clinical groups. Comparisons included (i) stroke subtypes versus healthy controls and (ii) ischemic stroke versus intracranial haemorrhage groups. Differentially regulated peptide sequences were subsequently subjected to sequence mapping and peptide locus analysis to determine regional clustering of endogenous peptides within precursor proteins. Peptide localization and sequence alignment were visualized in Python programming language. Furthermore, differentially regulated peptides were ranked based on their discriminatory performance using Mann–Whitney U testing and feature ranking approaches. Selected peptide candidates were subsequently used for development of an expression-guided decision tree classifier implemented in Python programming language. Following peptide selection, physicochemical properties of candidate peptides were calculated and integrated into downstream analyses as described in the subsequent section.

### Physicochemical characterization of differentially regulated peptides

Peptides identified as significantly upregulated or downregulated between intracerebral hemorrhage (ICH) and acute ischemic stroke (AIS) were extracted from the peptidomic dataset and analyzed separately. For each peptide sequence, a panel of physicochemical descriptors was calculated using a combination of custom Python scripts and established bioinformatics libraries. Custom sequence-derived metrics included the grand average of hydropathy (GRAVY) and aliphatic index. Additional physicochemical properties were calculated using the BioPython ProtParam module, including molecular weight, isoelectric point (pI), aromaticity, and instability index. Peptide bioactivity-related descriptors were further obtained using the modlAMP package, including charge, charge density, Boman index, hydrophobic ratio, and additional charge- and stability-associated features.

All calculated descriptors were assembled into feature matrices in which rows represented individual peptide sequences and columns represented physicochemical properties. Comparative analyses were subsequently performed between peptide groups originating from the AIS and ICH cohorts. For each numeric descriptor, differences between groups were quantified using mean shifts, Cohen’s d effect sizes, and two-sample Kolmogorov–Smirnov (KS) statistics to evaluate distributional separation. Descriptors were ranked according to their discriminatory power, and the highest-ranking features were selected for visualization. Distributional differences between groups were examined using overlaid normalized histograms generated with 15 bins, enabling assessment of shifts in feature distributions between AIS- and ICH-associated peptide populations. The resulting feature matrices were designed to provide a multidimensional physicochemical representation of disease-associated peptide repertoires and to facilitate subsequent biomarker discovery, statistical analysis, and machine-learning-based classification approaches.

### High-Throughput 96-Well Plate adaptation of the native serum peptidomics workflow

To enable scalable, high-throughput sample preparation, the benchtop native serum peptidomics protocol was adapted to a 96-well plate format. All steps were performed at 4°C using pre-cooled plates and equipment, unless otherwise stated.

Buffer preparation: The working solutions were prepared fresh on the day of each experiment using LC-MS grade water and reagents.

Sample processing: Serum samples were thawed on ice and clarified by centrifugation at 14,000 × g for 3 min at 4°C to remove particulate debris. A 100 µL aliquot of clarified serum was combined with 7× the volume of freshly prepared citrate-phosphate buffer in a pre-cooled 96-well deep-well plate. Samples were mixed by vigorous rocking on a platform rocker at maximum speed for 5 min at 4°C to dissociate endogenous peptide-protein interactions under native acidic conditions. The resulting mixture was diluted 1:1 (v/v) with 0.2% formic acid in LC-MS-grade water and clarified by centrifugation at 4,000 rpm for 5 min at 4°C.

Solid-phase extraction using Oasis HLB 96-well plate: Peptides purification were performed using an Oasis HLB 96-well SPE plate under positive pressure (20 psi) applied via a 96-well positive-pressure manifold. (i) Conditioning: each well was conditioned twice with 0.2% formic acid/methanol (v/v), applying gentle pressure using the left manifold knob (displaced a few mm). (ii) Equilibration: each well was equilibrated with 0.2% formic acid/water (v/v) under the same gentle pressure conditions. (iii) Sample loading: the clarified sample supernatant was loaded in two sequential additions, owing to the well capacity limit, applying only gentle pressure via the left manifold knob to maintain a controlled, low flow rate. (iv) Washing: each well was washed three times with water/5% methanol/0.2% formic acid (v/v) under gentle pressure. (v) Elution: the collection plate was replaced with a clean 96-well deep-well plate, and bound peptides were eluted with water/80% methanol/0.2% formic acid (v/v) per well.

Ultrafiltration: Peptide eluates were further purified by ultrafiltration using an Advance 96-well filter plate (Pall Corporation). Prior to use, the ultrafiltration membrane was pre-washed with 40% methanol per well under maximum positive pressure. Eluates from the SPE step were transferred to the pre-washed ultrafiltration plate in two sequential additions to avoid overfilling individual wells. The ultrafiltration flow-through, containing the enriched native serum peptide fraction, was collected into a glass-coated 96-well deep-well autosampler-compatible plate (Thermo Scientific SureSTART) The ultrafiltration membrane was kept wet throughout the process to prevent non-specific peptide adsorption and loss. Ultrafiltration was carried out for approximately 4 h under continuous positive pressure.

Sample storage and drying: Following ultrafiltration, samples were either stored directly at −80°C in the collection plate or dried overnight using a nitrogen evaporation system adapted for 96-well plates. Alternatively, samples were transferred to low-binding microcentrifuge tubes and dried by vacuum centrifugation (SpeedVac). Dried samples were stored at −80°C until mass spectrometric analysis. The entire 96-well workflow, from serum input to dried peptide extract, is completed within a single working day using less than 100 µl of serum per sample at a reagent cost of approximately €5–7 per sample, with no specialized liquid-handling robotics required.

## Data availability

The mass spectrometry intracellular peptidomics data have been deposited to the PRIDE repository.

## Authors contributions

S.K. Conceptualization, Methodology, Investigation, Supervision, Writing - Original Draft, Review & Editing, Funding Acquisition; J.F. Conceptualization, Methodology, Investigation, Writing - Original Draft, Review & Editing, Validation, Visualization; M.M. Methodology, Investigation, Visualization, Writing - Original Draft, Review & Editing; A.P. Investigation, Review & Editing; P.C. Investigation, Review & Editing, B.K. Investigation, Review & Editing; T.H. Funding Acquisition; N.M.T. Funding Acquisition.

## Acknowledgements

The authors acknowledge CI TASK, Gdańsk, Poland, the PLGrid Infrastructure, ACK Cyfronet AGH, and the Ares supercomputer for providing computational resources and technical support within grants no. plgneoantigen (PLG/2024/017749) and plgmsneoantigens (PLG/2024/017798). The authors are thankful to Prof. Piotr Trzonkowski and Dr. Mateusz Gliwiński, Gdańsk Medical University, for providing initial samples of Autoimmune disease.

