## Supplementary figures for "Novel native serum peptidomics workflow enables the discovery of circulating subtype-specific peptide biomarkers in acute ischemic and haemorrhagic stroke"

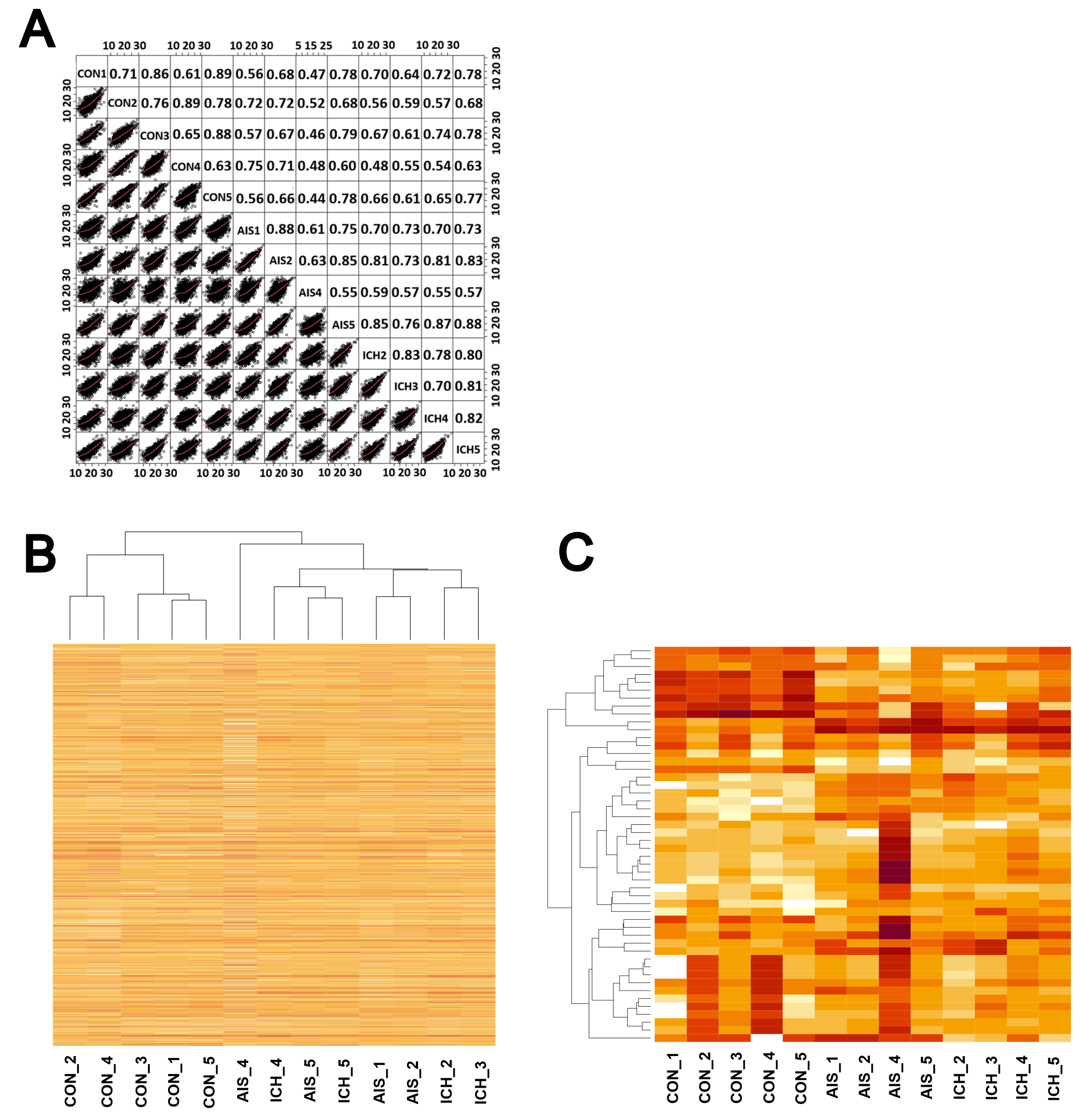

***Supplementary Figure 1.*** *Diagnostic plots of quantitative DIA serum/plasma peptidomics data. A) Scatter plots showing Pearson correlation coefficients (r-values) and regression between acute ischemic stroke (AIS), intercranial haemorrhagic stroke (ICH) and controls (CON). B) Heatmap with hierarchical clustering illustrating global serum peptidotype patterns across all samples. C) Filtered heatmap focusing on peptides with the highest fold-change standard deviation (STDEV cutoff > 3.4) enhancing stroke group separation.*

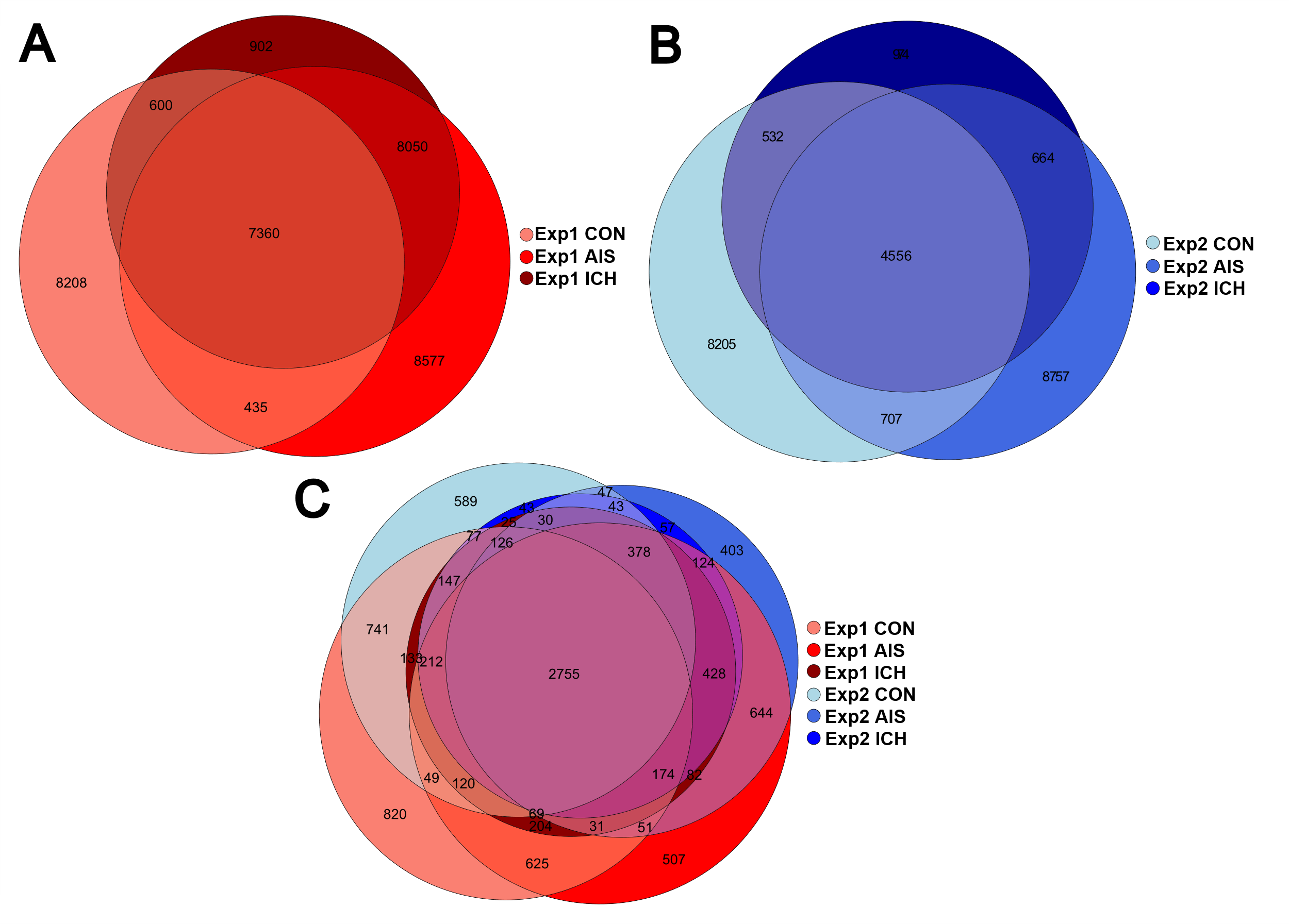

***Supplementary Figure 2.*** *Serum peptides overlap across experimental batches. Venn diagrams shoving the overlap of identified peptides among serum peptide between acute ischemic stroke (AIS), intracranial haemorrhagic stroke (ICH), and healthy controls (CON) from a single sample set processed in two independent batches of one sample set (Exp1 and Exp2). A) Peptide overlap between AIS, ICH and CON in Exp1, B) Peptide overlap between AIS, ICH and CON in Exp2. C) Peptide overlap between AIS, ICH and CON in both Exp1 and Exp2.*

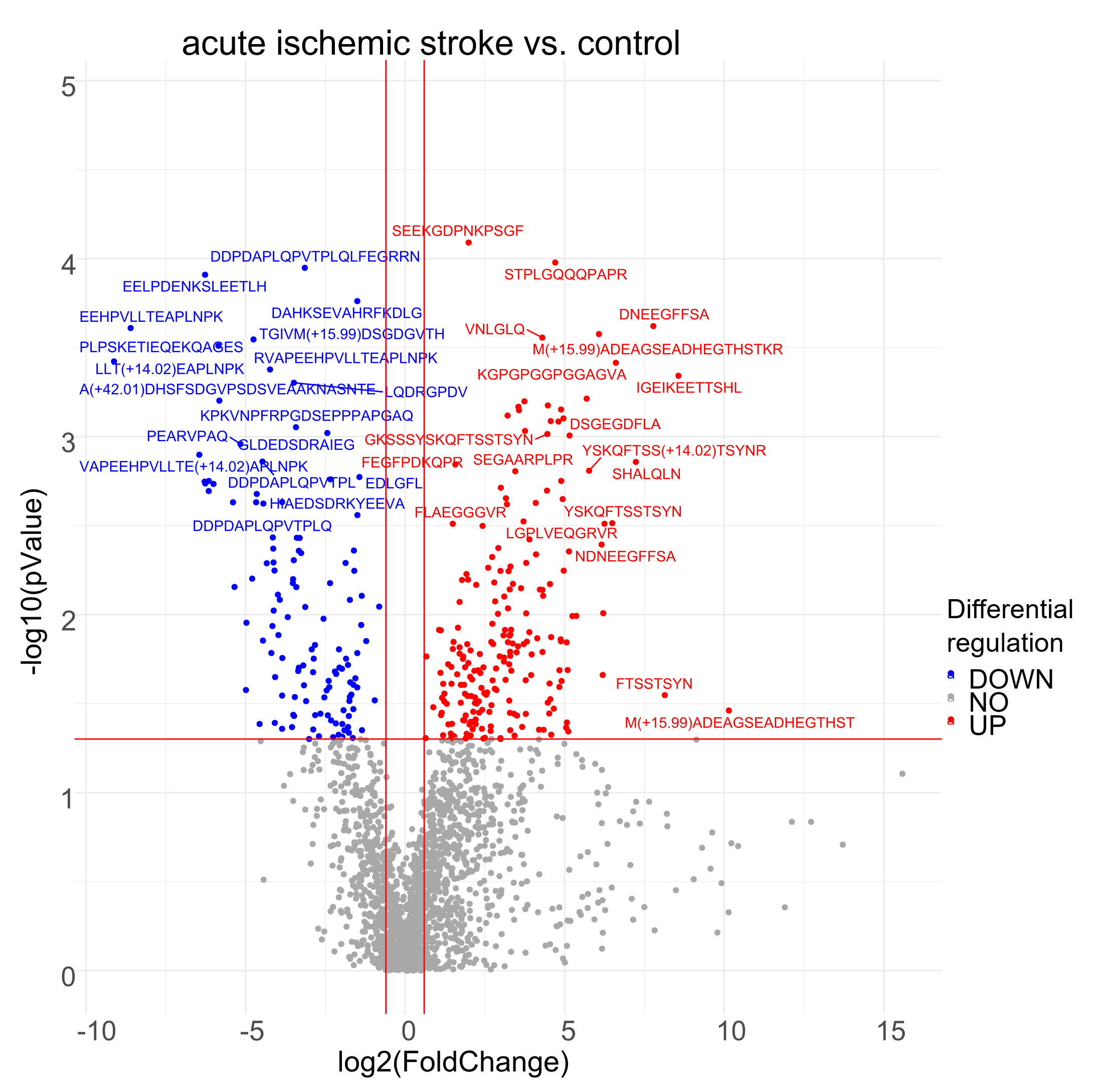

***Supplementary Figure 3.*** *Volcano plot comparing serum peptide levels between acute ischemic stroke (AIS) and controls. Peptides were considered significant at an adjusted P value ≤ 0.05 and a log_2_ fold-change ≥ 0.58 or ≤ −0.58.*

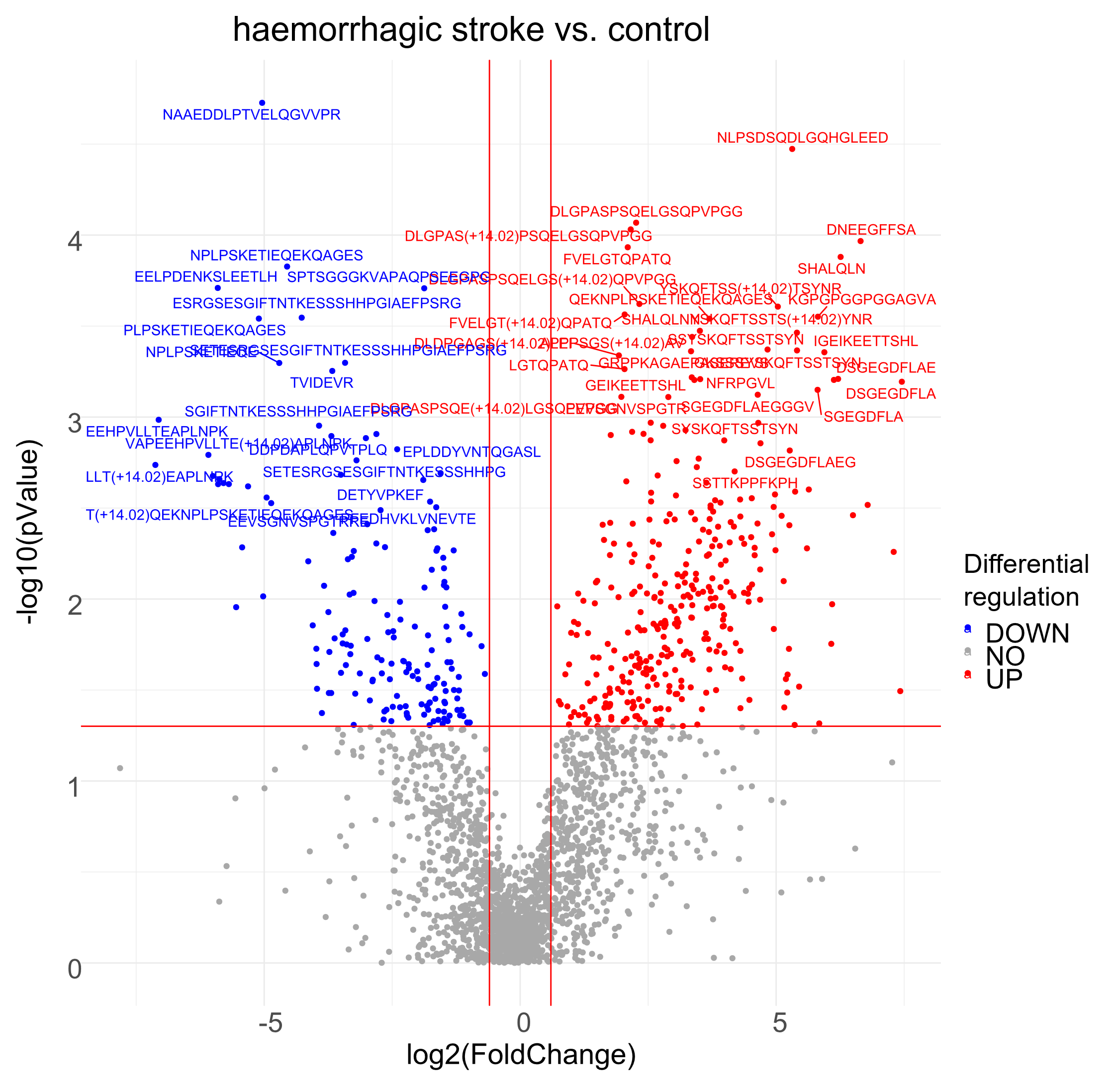

***Supplementary Figure 4.*** *Volcano plot comparing serum peptide levels between intercranial haemorrhagic stroke (ICH) and controls. Peptides were considered significant at an adjusted P value ≤ 0.05 and a log_2_ fold-change ≥ 0.58 or ≤ −0.58.*

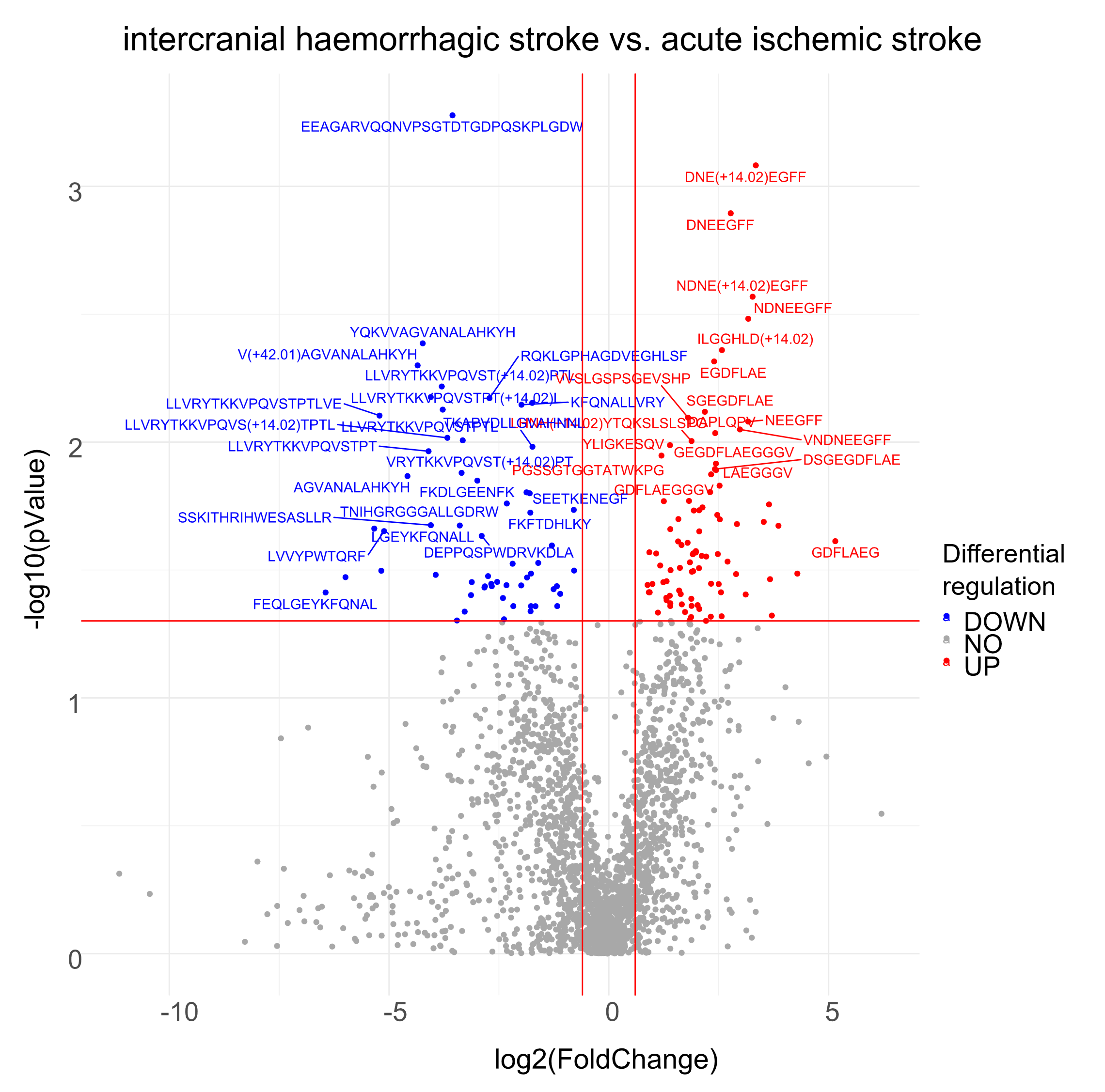

***Supplementary Figure 5.*** *Volcano plot comparing serum peptide levels between intercranial haemorrhagic stroke (ICH) and acute ischemic stroke (AIS). Peptides were considered significant at an adjusted P value ≤ 0.05 and a log_2_ fold-change ≥ 0.58 or ≤ −0.58.*

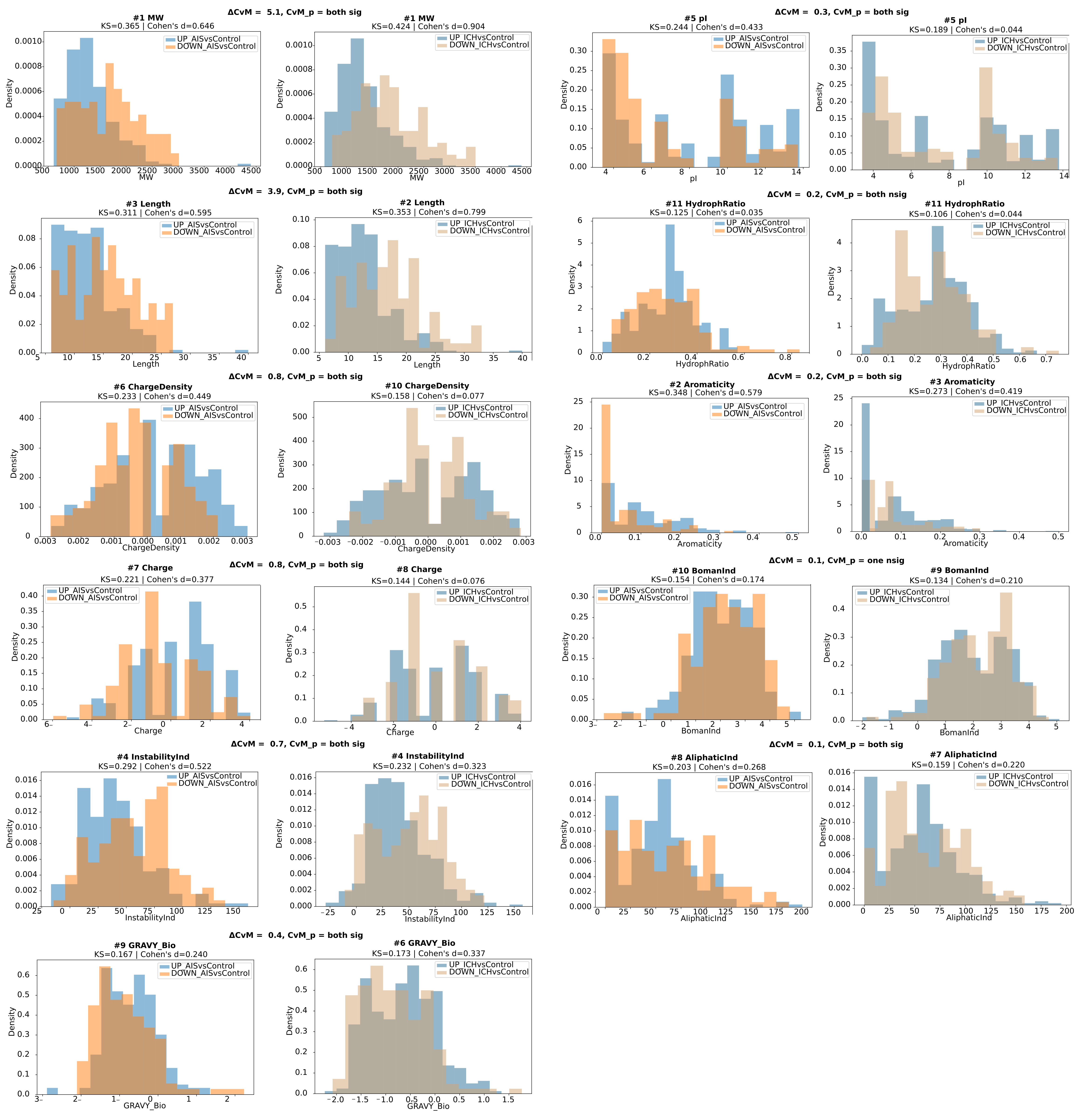

***Supplementary Figure 6. Distributional shifts in peptide physicochemical properties between stroke-associated and control-derived peptide populations.*** *Overlaid histograms showing the distributions of selected physicochemical descriptors for up- and down-regulated peptides identified in comparisons of acute ischemic stroke (AIS) and intracerebral haemorrhage (ICH) versus healthy controls. The magnitude of distributional differences between up- and down-regulated peptide populations was determined by Kolmogorov–Smirnov (KS) statistics and Cohen’s* d *effect sizes. Displayed descriptors include molecular weight (MW), aromaticity, peptide length, instability index, isoelectric point (pI), charge density, net charge, GRAVY index, and hydrophobic ratio. The differences in physicochemical property distributions between up- and down-regulated peptides in ICH versus controls and AIS versus controls were quantified using pairwise two-sample Cramér–von Mises (CvM) statistics, with differential distributional distances expressed as ΔCvM between conditions. Descriptors in the figure were ranked according to the absolute magnitude of ΔCvM values in descending order, thereby prioritizing descriptors exhibiting the greatest divergence in distribution comparisons. Histograms were generated from peptide-level physicochemical annotations derived from sequence-based descriptor analysis.*
